# Genomovar-level resolution reveals rapid pathotype switching and genomovar-specific disease potential in diarrheagenic *Escherichia coli* populations in northern Ecuador

**DOI:** 10.64898/2026.08.24.746700

**Authors:** Dorian J. Feistel, Kelsey J. Jesser, Karen Levy, Gabriel Trueba, Konstantinos T. Konstantinidis

**Affiliations:** Georgia Institute of Technology; University of Washington; Universidad San Francisco de Quito

## Abstract

Diarrheagenic *Escherichia coli* (DEC) pathotypes are commonly defined by molecular detection of discrete virulence genes, yet how quickly these diagnostic genes emerge and move among co-circulating lineages remain unclear. Here, we classified 248 whole-genome-sequenced *E. coli* isolates from the EcoZUR case-control study in northern Ecuador into intra-species genomovar units using the recently described 99.5% ANI threshold. This framework exposed cryptic population structure, revealing that single sequence types, representing identical multilocus sequence types (MLST), can harbor multiple distinct genomovars. Within individual genomovars, we observed a few cases of different pathotypes among isolates showing ∼99.7% ANI (and many such cases between genomovars). Coupled with synteny and phylogeny analyses that revealed pervasive incongruences between pathotype-diagnostic virulence genes and the core genome, these findings suggest recent horizontal gene transfer as the primary driver of pathotype evolution. Virulence gene profiling further revealed that accessory virulence repertoires are hierarchically structured by phylogroup across pathotypes, with genomovars assigned to phylogroups B2 and D exhibiting more conserved virulence architectures than those in phylogroup A and B1. Among DAEC isolates specifically, the B2- and D-associated genomovars showed elevated diarrhea-association rates relative to their phylogroup A counterparts. Rare virulence genes, including Type VI secretion systems, further distinguished diarrhea-associated from asymptomatic genomovars. These findings demonstrate that, although there seems to be within-lineage (phylogroup) conservation of virulence, pathotype identity is a labile state defined by horizontally acquired virulence genes at the genomovar level, and that the genomovar framework provides a biologically meaningful unit for linking intra-species diversity to pathogenic potential and outbreaks.

**Importance:** Efforts to diagnose diarrheal *Escherichia coli* infections depend on our ability to reliably identify which strain is dangerous, a task that for decades has rested on sorting strains into pathotypes defined by a few virulence genes. Whether these labels mark stable lineages or fleeting states is hard to judge with traditional typing methods such as Sequence Types (STs), which group together isolates with identical sequences in a handful of housekeeping loci. Using a recently defined genome-wide threshold (99.5% ANI), we resolved isolates into fine-scale genomovars and found that individual STs often conceal multiple distinct genomovars, some carrying conflicting virulence repertoires. At this resolution, the loci that define pathotypes are gained and lost far faster than the core genome diverges, and a genomovar’s genomic background shapes its association with disease. Genomovars therefore complement MLST with the resolution needed to interpret genomic surveillance data and to build robust diagnostic and public-health frameworks.

## Introduction

Diarrheal diseases remain among the leading causes of morbidity and mortality worldwide, disproportionately affecting children under five years of age in low- and middle-income countries, where inadequate water, sanitation, and hygiene (WASH) infrastructure perpetuates transmission. Among the diverse etiological agents of infectious diarrhea, diarrheagenic *Escherichia coli* (DEC) pathotypes^1^—including enterotoxigenic *E. coli* (ETEC), enteroaggregative *E. coli* (EAEC), enteropathogenic *E. coli* (EPEC), enteroinvasive *E. coli* (EIEC), diffusely adherent *E. coli* (DAEC), and enterohaemorrhagic *E. coli* (EHEC)—are collectively recognized as major contributors to the global diarrheal disease burden^2–4^. These pathotypes are distinguished by their characteristic virulence mechanisms: ETEC delivers secretory enterotoxins into the epithelial cells of the small intestine^5^, EAEC forms adherent biofilms on the intestinal epithelium^6^, EPEC induces attaching and effacing lesions^7^, EIEC invades the colonic mucosa^8^, and DAEC elicits elongation of microvilli through diffuse adherence mediated by the Afa/Dr family of adhesins^9^. Despite this mechanistic understanding, the genomic architecture underpinning the emergence, diversification, and pathogenic potential of co-circulating DEC lineages varies considerably in how well it is characterized across pathotypes—comparatively well-defined for some (e.g., EIEC) but poorly understood for others (e.g., DAEC)—and remains especially underexplored at intra-species resolution within individual epidemiological cohorts.

The EcoZUR (*E. coli* en Zonas Urbanas y Rurales) study was a case-control study investigating the etiology of diarrheal disease in communities spanning an urban–rural gradient in northern Ecuador, conducted over an 18-month period from 2014-2015. The study has provided a uniquely comprehensive framework for examining DEC epidemiology and genomics within a geographically defined human population^10–12^ and has yielded findings that collectively frame the present work. First, Montero *et al.*^13^ characterized the distribution of DEC pathotypes across the urban–rural gradient and demonstrated that DAEC, the most prevalent pathotype overall, was more frequently associated with disease in urban settings and exhibited extensive genetic diversity among diarrhea-associated isolates, with no evidence that disease associations were driven by a single virulent clone(s). Second, Peña-González *et al.*^14^ applied a novel metagenomic framework demonstrating that DAEC was the likely causative agent of diarrhea in approximately half of culture-positive samples, that DAEC infections were uniquely accompanied by co-elution of large amounts of human DNA and significant shifts in gut microbiome composition, and that pathotype-specific signatures in the diseased gut microbiome could distinguish DAEC from ETEC infections, which were subsequently further validated with more detailed metagenomic analysis^15^. Third, Rothstein *et al.*^16^ employed phylodynamic and phylogeographic methods on DEC genomes from EcoZUR and found minimal phylogenomic structuring by site, pathotype, or clinical status (diarrhea vs asymptomatic), revealing instead extensive mixing of DEC pathotypes across the study sites, with migration rates from urban towards rural populations estimated to be 6.7-fold higher than in the reverse direction. Notably, all these studies converged on the observation that DEC—and the DAEC pathotype in particular—circulate as genetically diverse, polyphyletic assemblages whose pathogenic potential cannot be readily explained by conventional pathotype classification or clonal structure alone, thereby motivating finer-resolution genomic analyses to identify the determinants of virulence among co-circulating strains.

The convergence of extensive genotype mixing, polyphyletic pathotype distributions, and variable disease associations observed in the EcoZUR study raises several interrelated questions that conventional classification schemes have been unable to resolve. First, if pathotype designations are conferred by discrete, horizontally acquired virulence genes, how rapidly can these genes move among co-circulating lineages? Second, does the phylogenetic or lineage background modulate disease potential independently of pathotype? Third, given the 6.7-fold asymmetry in urban-to-rural migration rates documented by Rothstein *et al.*^16^, are the genomic determinants of virulence plasticity disseminated broadly across the transmission landscape, or do they remain confined to specific lineages?

To address these questions, we report a comprehensive, intra-species-resolution genomic characterization of 248 *E. coli* isolates from the EcoZUR study. We applied the 99.5% ANI framework^17^ to resolve isolates into genomic units (i.e. genomovars) and then integrated pangenomic characterization, core genome phylogenomics, pathotype assignment, and comprehensive virulence factor profiling to examine population structure, horizontal gene transfer dynamics, and lineage-associated variation in pathogenic potential. By using tanglegrams to compare diagnostic virulence gene similarity to core genome phylogeny, we traced discrete lateral transfer events to assess the timescale and directionality of pathotype-defining gene mobility. We further evaluated whether phylogroup- and genomovar-level differences in accessory virulence gene content corresponded to isolation from human hosts with and without acute diarrhea symptoms, and whether rare gene signatures can distinguish genomovars that are associated with diarrheal cases from those more commonly associated with asymptomatic carriage. Together, these analyses provide a genomovar-resolution portrait of DEC population dynamics in an endemic setting and offer a framework for understanding how intra-species genomic diversity shapes the emergence and dissemination of virulent *E. coli* lineages.

## Methods

### DNA extraction, sequencing, and data availability

The EcoZUR project^18^ collected fecal samples from individuals with diarrhea and age- and location-matched controls living in four regions spanning an urban to rural gradient in Ecuador (Quito, Esmeraldas, Borbón, and several rural villages located along the Cayapas, Santiago, and Onzole Rivers). Individuals of all ages were recruited between April 2014-September 2015 from Ecuadorian Ministry of Health hospitals and/or clinics in each location. *E. coli* isolation methods are described elsewhere^14^. DNA from *E. coli* colonies that tested positive for a virulence factor of interest was extracted using the Wizard Genomic DNA Purification kit (Promega) and the extract purity and concentration was estimated using a NanoDrop spectrophotometer (Thermo Scientific) and the Qubit 2.0 dsDNA high-sensitivity assay (Invitrogen, Carlsbad, CA). DNA sequencing libraries were prepared using the Illumina Nextera XT DNA library preparation kit (Illumina) according to manufacturer’s instruction. After this, libraries were run on a High Sensitivity DNA chip using the Bioanalyzer 2100 instrument (Agilent) to determine library insert sizes. An equimolar mixture of the libraries (final loading concentration of 10 pM) was sequenced on an Illumina MiSeq instrument (School of Biological Sciences, Georgia Institute of Technology), using a MiSEQ reagent v2 kit for 500 cycles (2 x 250 bp paired end run). Adapter trimming and demultiplexing of sequenced samples was carried out using the MiSEQ control software v2.4.0.4. In total 316 *E. coli* isolates tested positive for any of the nine PCR assays and from those, 279 were successfully sequenced. The complete set of strains sequenced in this study has been deposited in the NCBI Sequence Read Archive (SRA) under BioProject ID PRJNA486009.

### Read quality control and de novo assembly

Raw sequencing reads from *E. coli* isolates were quality-assessed using FastQC (https://github.com/s-andrews/FastQC). Adapter trimming and quality filtering were performed using fastp^19^ with automatic paired-end adapter detection (--detect_adapter_for_pe). Bases with a Phred quality score below Q20 were trimmed and reads shorter than 50 bp after trimming were discarded. Taxonomic classification was conducted using Kraken2^20^ with species-level abundance estimation via Bracken^21^ and visualized with custom scripts (https://github.com/rotheconrad/Kraken-Bracken-plot). De novo genome assembly was performed using SPAdes^22^ v3.13.0 with the --careful flag to minimize short insertions, deletions, and substitution errors. Assembled contigs were filtered to retain only those ≥500 bp with k-mer coverage ≥2. Assembly quality was evaluated using CheckM^23^ with only genome assemblies with >90% completeness and <5% contamination retained for downstream comparative genomic analyses.

### Pathotype assignment of E. coli isolate genomes

*E. coli* draft assemblies were assigned a pathotype based on the presence of pathotype diagnostic genes (Table 1) using a previously described read mapping-based approach^14^. Briefly, quality-filtered reads were aligned to pathotype-specific virulence gene sequences using BLAST^24^ with minimum thresholds of 90% sequence identity and 70% coverage. Best-matching alignments were retained using BlastTab.best_hit_sorted.pl^25^ and average sequence depth was calculated using BlastTab.seqdepth_ZIP.pl^25^. Pathotype diagnostic genes with average sequencing depth ≥1X were scored as present. Isolates were classified according to pathotype diagnostic gene combinations defined in Table 1; those harboring genes associated with multiple pathotypes were designated as hybrids^26^. Given the observed discordance between PCR- and genome-based pathotype assignments (61/247 isolates), with most discordant isolates (40/61) lacking detectable pathotype diagnostic genes at the defined thresholds, pathotype assignments used in downstream analyses were based on the in silico read-mapping approach, which provided a standardized genome-wide assessment of pathotype diagnostic genes

**Table 1:**
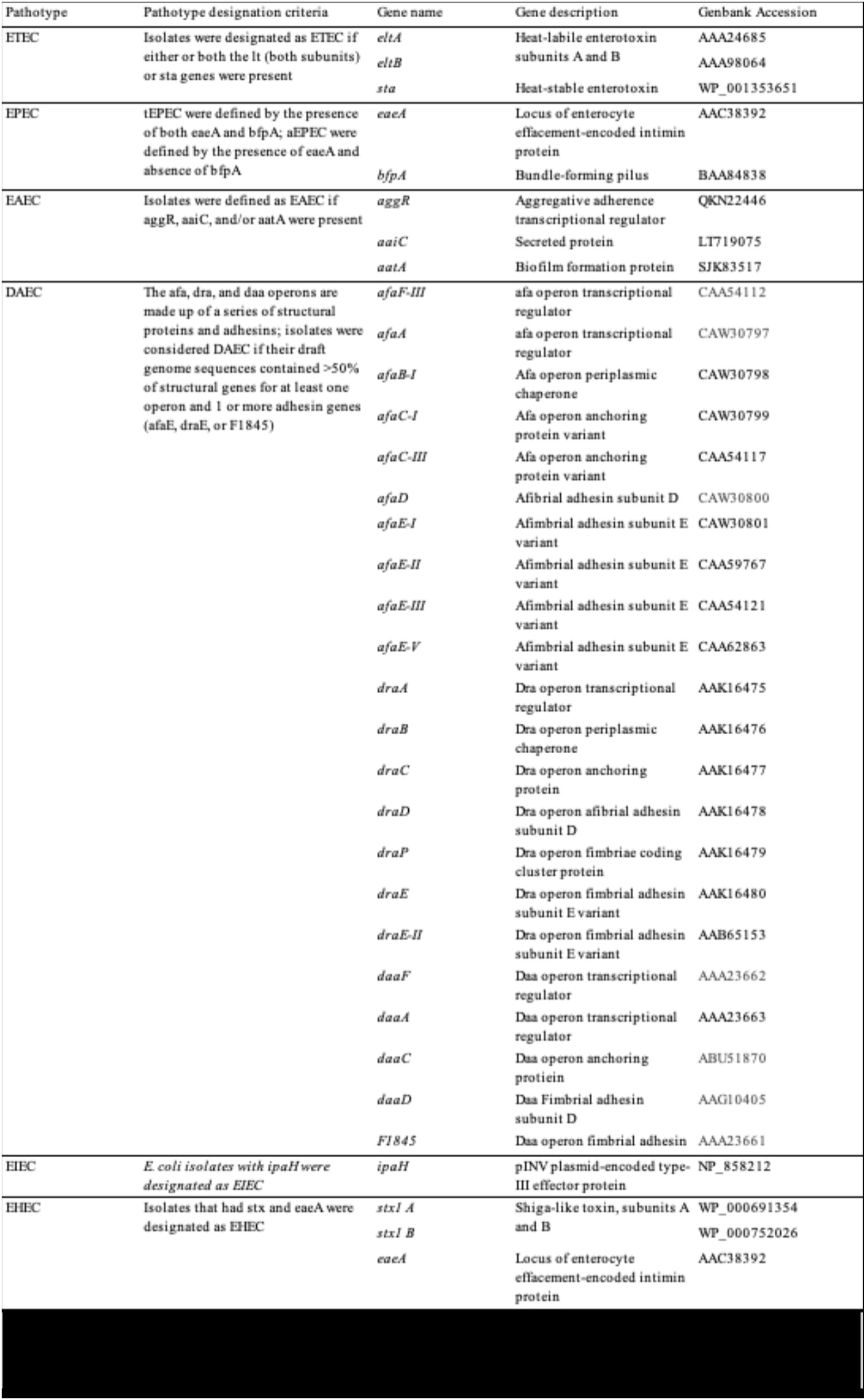
Diagnostic genes use for pathotyping assignment.

### Average nucleotide identity and genomovar delineation

Genome ANI was calculated using FastANI^27^ v1.33 with default settings. Scatterplots with histograms visualizing the relationship between ANI and the shared genome proportion (calculated as the number of mapped bidirectional fragments divided by the total number of fragments) were created with custom scripts using Python v3.12.12. Self-matching comparisons were excluded and only a single ANI value was retained for each pairwise comparison, corresponding to the larger genome in the dataset. A third-degree polynomial regression was fitted to the ANI versus shared genome proportion data, and goodness of fit was assessed using the coefficient of determination (*R*^2^). A symmetric distance matrix (calculated as 1 – ANI/100) was subjected to hierarchical clustering using the “linkage” function from *scipy.cluster.hierarchy* (SciPy v1.9.3^28^) with default settings except using a complete-linkage method. Genomovars were then assigned by partitioning the hierarchical tree into flat clusters using the “fcluster” function from *scipy.cluster.hierarchy* with a distance threshold (t) of 0.005 (which corresponds to 99.5% ANI). The cophenetic correlation coefficient, measuring similarity between original pairwise distances and dendrogram-derived cophenetic distances, was computed using the “cophenet” function from *scipy.cluster.hierarchy*.

### In silico inference of sequence types and phylogenetic classification

Sequence Types (ST) were assigned using the Warwick (Achtman) scheme^29^, which classifies isolates based on allelic variation across seven housekeeping loci (*adk, fumC, gyrB, icd, mdh, purA,* and *recA*). Isolate genomes were assigned STs *in silico* using mlst (http://github.com/tseemann/mlst) v2.22.0 with the “ecoli_achtman_4” database. ANI distribution plots for discrete STs were constructed using similar methods as described above (refer to *Average Nucleotide Identity and Genomovar Delineation*). For classification of ANI-based ST concordance, pairwise genome comparisons were labeled as true positives (TP; same ST and ANI ≥ 99.5%), true negatives (TN; different ST and ANI < 99.5%), false positives (FP; same ST and ANI < 99.5%), or false negatives (FN; different ST and ANI ≥ 99.5%). For each discrete ST, metrics were calculated in a one-vs-rest framework using all comparisons that included that ST (i.e., ST-to-ST and ST-to-non-ST pairs), and precision (TP / (TP + FP)), recall (TP / (TP + FN)), F1 (2 × (Precision × Recall) / (Precision + Recall)), and accuracy ((TP + TN) / (TP + TN + FP + FN)) were computed from the corresponding TP/TN/FP/FN counts. Overall performance was summarized using macro averages (unweighted mean of per-ST metrics), micro averages (metrics from pooled global counts across all included comparisons), and weighted averages (per-ST metrics weighted by ST support/datapoints). A minimum-genome threshold was applied where only STs with three or more genomes were included in the overall aggregate summaries.

Clermont phylogroup membership was assigned *in silico* using ClermonTyping^30^. To validate and, when necessary, refine these assignments, we leveraged genome-wide relatedness inferred from the symmetric matrix of pairwise ANI values. We visualized this matrix as a hierarchical clustering map and annotated the clustered genomes with their ClermonTyping phylogroup calls (data not shown). Because primer-binding variation and horizontal gene transfer can occasionally bias PCR-derived phylogrouping^30^, we used concordance between ANI-based clustering and phylogroup labels as a consistency check. When a genome’s ClermonTyping assignment was inconsistent with its placement in the ANI cluster map, we reassigned the phylogroup based on its closest genomic neighbors based on ANI. Thus, genomes lacking a supported phylogroup call were assigned to the phylogroup shared by genomes with which they clustered at approximately 98.5% ANI. Conversely, when a genome’s inferred phylogroup conflicted with the predominant phylogroup of its genomovar, we reassigned it to match the phylogroup most common within that genomovar.

### Core gene phylogenetic reconstruction

Coding sequences (CDS) were predicted using Prodigal^31^ and sequences shorter than 300 bp were discarded. Predicted CDS were clustered using CD-HIT^32^ with the following parameters: “-c 0.9 -n 8 -G 0 -g 1 -aS 0.7 -M 10000 -d 0 -T 10.” Paralogs were removed within each cluster by keeping only the sequence with the highest percent sequence identity to the cluster representative. Conserved core genes, defined as orthologous genes shared between >99% of isolates, were aligned using MAFFT^33^ with default parameters. The resulting sequence alignments were concatenated into a single multiple sequence alignment using the *Aln.cat.rb*^25^ script with the “--remove-invariable” flag, creating a SNP alignment. The phylogenetic tree was constructed using RAxML^34^ v8.2.12, employing the GTRGAMMA substitution model and 1000 bootstrap replicates and visualized with metadata using iTol^35^.

### Defining and detecting recent horizontal gene transfer events

Genome assemblies (contigs) were annotated with Bakta^36^ to predict gene functions and determine gene synteny. DAEC genomes containing *afa/dr* operon CDSs corresponding to the conserved A/B/C/D/P/E gene components in a collinear arrangement on a single contig were extracted. *afa/dr* genes were grouped by their conserved component designations (A, B, C, D, P, and E) to combine corresponding genes from *afa*, *dra*, and *daa* operon variants, and multiple sequence alignment was performed using MAFFT^33^. An approximate-maximum likelihood *afa/dr* operon tree was generated using FastTree^37^ v2.1.11 using “-nt -gtr” flags. The tanglegram was manually curated by aligning the core gene phylogeny to the *afa/dr* operon tree and annotated with the genomovar and phylogroup assignments using iTol^35^. Tanglegrams for additional pathotype-diagnostic genes listed in Table 1 were generated using the same workflow described above, except that phylogenies were inferred from individual gene alignments. We inferred horizontal gene transfer (HGT) of the *afa/dr* operon when isolates showed incongruent placement between the core genome and operon phylogenies. The same approach was used to detect HGT events for the diagnostic genes of other pathotypes listed in Table 1 (see tanglegrams in Supplementary Figure 1). In addition, we considered additional evidence to support HGT: i) when observed operon substitutions deviated from expectations derived from genome-wide ANI—either among variants within the same genomovar or between closely related genomovars within the intra-species ANI gap, and ii) when distantly related genomovars from distinct phylogroups shared ≤5 substitutions across the complete operon sequence.

To enable comparative, locus-centered analyses, predicted CDS were compared in an all-versus-all manner using the *aai.rb* script^25^, which retains a single reciprocal best matching gene to define putative orthologous correspondences across genomes. In cases where multiple reciprocal matching genes with similar identities (paralogs) were identified, one was chosen at random. For percent sequence identity-based visualization, the genome that had the longest contig harboring the gene(s) of interest was selected as a reference. Reference contigs were ordered by descending length, and all matched genes from other genomes were projected onto the reference coordinate system based on the relative position of their reciprocal best match along the corresponding reference contig. Comparative plots were used to display genomic position along the reference (x-axis) versus nucleotide sequence identity of the reciprocal best match (y-axis) and additionally incorporate per-genome metadata to contextualize variation consistent with horizontal gene transfer and related evolutionary events. Genes were annotated with functional categories using Bakta-derived metadata or classified as “pathotype-associated” genes, which included the pathotype-diagnostic genes listed in Table 1. Bakta annotations were grouped into five functional categories: virulence factors, mobile genetic elements (insertion sequences, transposases, integrases), structural/regulatory elements (gap, oriC, oriT, sorf, crispr), functional RNA elements (ncRNA, ncRNA-region, rRNA, tmRNA, tRNA), and hypothetical proteins. Genes lacking functional annotation from either source were assigned to pangenome categories based on their prevalence across all *E. coli* genomes sharing >95% ANI: conserved (>99%), core (≥90%), accessory (<90%), or rare (≤10%). For *afa/dr* operon synteny analysis, DAEC genomes in which all operon genes were located on a single contig were selected, and genes within a 15-gene upstream and downstream window of the *afaC* gene were extracted. Gene positions were plotted relative to *afaC* as the target point, with arrows indicating transcriptional direction, to identify flanking mobile genetic elements (insertion sequences, transposases, and integrases) and characterize variation in the genomic context surrounding the operon across lineages. Contigs harboring the *afa/dr* operon were additionally classified as chromosomal or plasmid in origin using PlaScope^38^.

### Virulence factor detection

The full Virulence Factors Database^39^ (VFDB) of known bacterial virulence factors was downloaded on 31 May 2022 and used as the reference for virulence factor identification in our isolate collection. Genomes sharing greater than 95% ANI were used in this analysis. Predicted CDSs were queried against VFDB using BLAST^24^ and matches were retained using ≥90% nucleotide sequence identity and ≥70% sequencing breadth of coverage (defined as alignment length divided by subject length). For downstream curation, retained matches were collapsed across allelic variants and harmonized to standard accepted virulence factor gene names based on VFDB nomenclature to generate a non-redundant presence/absence matrix. Genes belonging to the *afa*, *dra*, and *daa* operons associated with DAEC were collapsed under a unified nomenclature as they share structural and functional similarities^9,40,41^.

Virulence factor frequencies were summarized as the proportion of genomes in the dataset containing each gene and visualized as a histogram with an overlaid cumulative frequency curve. VF prevalence categories were defined using dataset-specific thresholds selected by visual inspection of inflection points in the cumulative frequency distribution, with genes detected in ≤15% of genomes classified as rare VFs, genes detected in ≥90% of genomes classified as core VFs, and intermediate-frequency genes classified as accessory VFs. Pathotype-specific virulence factors were identified by integrating diagnostic genes from Table 1 with additional candidates curated from peer-reviewed literature for the pathotypes detected in this study (Supplementary Table 2), prioritizing markers characteristic of individual or closely related pathotypes.^3,42,43^. Pathotype-specific genes were subsequently clustered using the *seaborn*^44^ “clustermap” function. Functional assignments for each virulence factor were taken from VFDB-associated metadata.

### Accessory and rare virulence factor analyses and statistical testing

A presence-absence matrix of accessory VFs (those with intermediate prevalence between >15% and <90% of genomes) was constructed across genomovars containing at least three members. Pairwise distances between genomes were calculated using the *<u>beta_diversity</u>* function from the *skbio.diversity* Python library with the Dice-Sørensen dissimilarity metric (<u>metric=’dice’</u>) applied to the binary accessory gene matrix, generating a distance matrix that quantifies genomic dissimilarity based on shared and unique accessory VF content. This distance matrix served as input for downstream dimensionality reduction and statistical analyses.

Permutational multivariate analysis of variance^45^ (PERMANOVA) was conducted on the Dice-Sørensen distance matrix to test for significant associations between accessory genome composition and both phylogroup classification and clinical response categories (diarrhea versus asymptomatic). Homogeneity of multivariate dispersion among groups was assessed using permutational analysis of multivariate dispersions (PERMDISP). PERMANOVA and PERMDISP were performed using the *permanova* and *permdisp* functions with 999 permutations from *skbio.stats.distance*^46^. Pairwise comparisons between phylogroups were conducted for both tests and resulting p-values were corrected for multiple comparisons using the Benjamini-Hochberg false discovery rate (FDR) method. To assess the correlation between accessory genome composition and genomic relatedness, a Mantel^47^ test was performed between the Dice-Sørensen distance matrix and the ANI distance (see above) using the “mantel” function from the *skbio.stats.distance* with 999 permutations and a Pearson correlation coefficient.

To visualize patterns of accessory genome variation, Uniform Manifold Approximation and Projection^48^ (UMAP) was performed on the Dice-Sørensen distance matrix setting the minimum embedding distance parameter to 0.35 and the local neighborhood size parameter set to 50. Genomes were plotted in two-dimensional UMAP space, with points colored by phylogroup assignment and shaped according to genomovar cluster membership to evaluate concordance between accessory genome architecture and phylogenetic classification using a custom Python script. To characterize the accessory virulence gene repertoire of each genomovar, genes were classified into functional categories using VFDB annotations, and the relative proportion of each category was calculated per genomovar. Proportions were visualized as stacked bar plots grouped by phylogroup to compare virulence factor functional composition across lineages.

The diarrhea case percent for each phylogroup was calculated as the number of diarrhea-associated genomes divided by the total number of sampled genomes and multiplied by 100 for the overall (aggregate) and within-DAEC-specific pathotype subset. An adjusted disease-association percentage with Wilson^49,50^ 95% confidence intervals was computed to account for uneven sampling, and the results were visualized as a horizontal point-and-whisker plot. Pairwise differences in disease-association rates between phylogroups were assessed using Fisher’s^51^ exact test on 2×2 contingency tables of diarrhea versus asymptomatic counts.

For comparative analysis of rare gene distributions between representative genomovars, the union of rare genes present in the selected genomovars was identified. A binary presence-absence heatmap was generated with genomes ordered along the x-axis according to their phylogenetic position in the core genome tree and rare genes displayed along the y-axis. The heatmap was annotated with phylogroup assignment, pathotype classification, virulence factor functional categories, and clinical outcome status for each genome to facilitate interpretation of rare gene distribution patterns in relation to phylogenetic and phenotypic characteristics.

## Results

### Genome sequencing and quality assessment of diarrheagenic E. coli isolates from northern Ecuador

Of the 271 diarrheagenic *E. coli* strains sequenced, 24 isolates showed a substantial signal (10 to 50% relative abundance) of genomic contamination with non-*Escherichia* genera or did not meet our threshold criteria of completeness (>90%) and contamination (<5%) as estimated by CheckM^23^. We concluded that these isolate genomes were most likely contaminated, and they were discarded from further analysis. The remaining 248 isolates were determined to be of high-draft quality following *de novo* assembly (mean completeness = 99.9%, SD = 0.61%; mean contamination = 0.4%, SD = 0.67%) and were further used to study the population structure of *E. coli* circulating in Northern Ecuador. These draft genomes consisted of 200.6 contigs on average (SD = 102.6), with an average G+C% content of 50.6% and an estimated genome size of 5.1Mbp (SD = 0.25Mbp; minimum = 4.39Mb; maximum = 6.19Mbp), and were predicted to contain an average of 4289.2 (SD = 216.1) putative protein-coding gene sequences with an average length of 1003.2bp (SD = 633.4bp). In addition, the average sequencing depth of coverage of the assembled genomes was 37.2X (SD = 10.1X; minimum = 10.58X; maximum = 70.53X).

### Genomovar delineation of EcoZUR diarrheagenic E. coli isolates reveals cryptic intra-species genomic structure within and between Sequence Types

Nearly all isolate genomes (247/248) exceeded the threshold for species delineation, with pairwise ANI values no lower than 96.3%; a single isolate (B37-6) fell below the species boundary (mean pairwise ANI = 92.8% ± 0.07% SD). We observed a discontinuity in our dataset mostly between 99.4–99.9% ANI (Figure 1) and adopted the genomovar framework by applying the 99.5% ANI midpoint as a threshold to hierarchical cluster the 248 EcoZUR *E. coli* isolates into genomovars. This approach resolved 50 multi-genome genomovars: 35 genomovars containing ≥3 isolates and 15 containing genome pairs, with 51 isolate genomes being assigned to a singleton genomovar (cophenetic correlation coefficient = 0.98).

**Figure 1:**
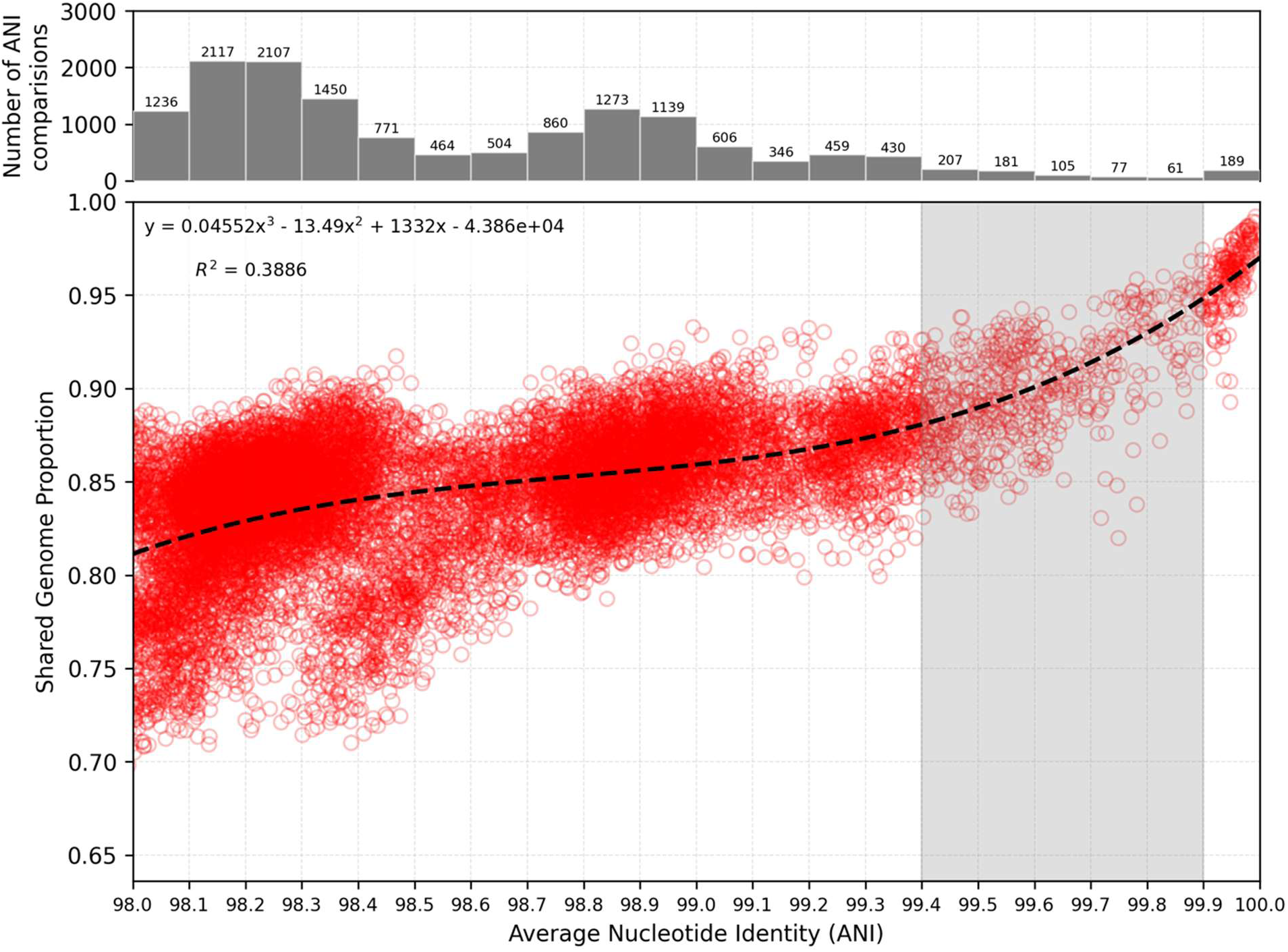
Genomic similarity among EcoZUR pathogenic *E. coli* isolates, measured by the shared genome proportion as a function of average nucleotide identity. Each data point corresponds to a pairwise ANI comparison and shared genome proportion between two genomes. The dashed black trendline represents a cubic polynomial regression fit to the data, capturing the overall relationship between ANI and genome sharing across the collection. The histogram (top panel) displays the frequency distribution of pairwise ANI values, showing the number of genome comparisons within each ANI interval. The shaded gray region (99.4–99.9% ANI) highlights a notable gap in the ANI distribution, with a noticeable scarcity of genome pairwise comparisons in this range—the frequency drops sharply from roughly 99.3% bin and remains low until 99.9%, then increases approximately three-fold in the 99.9–100% range.

Overall, there was a high degree of diversity in the STs for EcoZUR *E. coli* isolates, with >99% of the isolates assignable to known STs. A total of 93 nonredundant STs were identified, among which 26 contained three or more genomes, 11 were paired, and 56 were singletons. The largest STs included ST10 (n=30), ST38 (n=15), ST131 (n=15), ST4 (n=9), and ST270 (n=8). with most of these being well documented in the literature previously. For instance, the ability of these STs to cause disease and/or harbor antibiotic resistant genes has been described previously^52–60^. With respect to ST-10, the distribution was similar with previous studies in that it is one of the most frequently found STs in human fecal and food samples^61^.

For the four most abundant STs detected (ST-10, ST-131, ST-38, and ST-4), we observed that all exhibited a gap within the 99.2–99.8% ANI range, although the depth and breadth of this discontinuity varied by ST lineage (Figure 2). Establishing the previously defined 99.5% ANI midpoint^17^ as a boundary for genomovar delineation prior to ST assignments showed that pairwise comparisons frequently breached this boundary. For example, ST-matching genome pairs falling below the 99.5% ANI midpoint were common for ST-10 (precision = 0.228), ST-131 (0.457), and ST-38 (0.629), all well below the macro-averaged precision of 0.849, while non-matching ST pairs sharing ≥99.5% ANI were prevalent (although less common) for ST-4 (recall = 0.522) and ST-10 (0.702), both below the macro-averaged recall of 0.843. Taken together, these patterns expose a cryptic intra-species genomic structure that delineates distinct evolutionary lineages within STs (i.e. genomovars). These cryptic lineages—invisible to conventional MLST classification—demonstrate that sequence types do not always reliably delineate natural population boundaries at intra-species resolution, underscoring the need for genomovar-level analysis to complement, rather than replace, the well-established utility of MLST in epidemiological surveillance and lineage tracking. For subsequent results, we have designated each genomovar with a ‘g’ prefix followed by an arbitrary integer (e.g., g1, g2, g3) and leverage this classification to examine population structure, temporal dynamics, and phylogenetic relationships among our isolates, with particular emphasis on identifying genomovars that harbor distinct virulence gene profiles, evidence of horizontal gene transfer shaping pathotype diversity, and elevated pathogenic potential.

**Figure 2:**
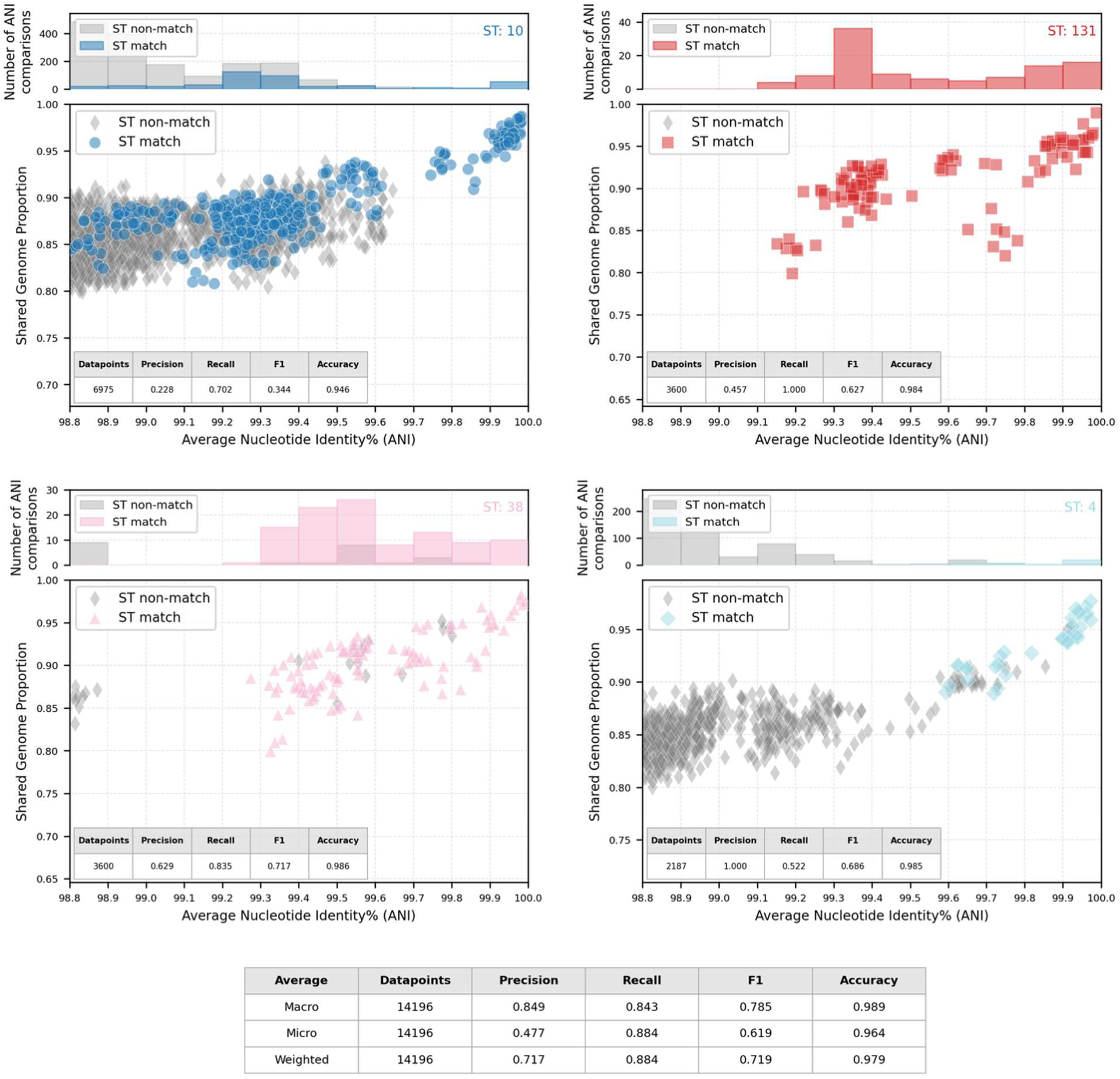
Average nucleotide identity (ANI) distribution for EcoZUR pathogenic *E. coli* genomes by sequence type (ST). Each of four panels focuses on a single ST and displays pairwise ANI versus shared genome proportion for all genome pairs in which at least one genome belongs to the focal ST. Within each panel, data points are classified as ST-matching (both genomes assigned to the focal ST) or ST non-matching (only one genome assigned to the focal ST). Histograms above each scatter plot show the ANI distribution for the corresponding pair category. Inset tables within each panel report classification metrics specific to that ST, while the table below summarizes macro-, micro-, and weighted-average metrics across all STs represented by three or more genomes (n = 26 STs). The analysis reveals that a substantial proportion of ST-matching genome pairs for ST-10 fall below the 99.5% ANI midpoint used as a threshold to delineate genomovars, indicating that genomes assigned to the same ST can span considerable genomic divergence despite their conserved core loci used in MLST. Conversely, some ST-non-matching pairs exceed 99.5% ANI (such as ST-38 and ST-4), demonstrating that genomes from different STs can be nearly indistinguishable at the whole-genome level despite differences in at least one of their seven housekeeping genes used for multi-locus sequence type (MLST) assignment. The breadth and depth of the upper ANI gap vary across STs, potentially reflecting differences in sampling depth, isolation biases, or intrinsic rates of sequence divergence within each ST.

### Pangenomic structure of EcoZUR diarrheagenic E. coli isolates reveal pathotype discordance within genomovars

We found that 208 (83.9%) isolate genomes harbored one or more pathotype-diagnostic genes meeting our assignment thresholds, while 40 (16.1%) isolates lacked sufficient markers and were classified as ‘Unknown’. Interestingly, five isolates carried diagnostic genes from both EAEC and ETEC pathotypes and were designated EAEC/ETEC hybrids. The pathotype distribution was: DAEC (n = 85), EAEC (n = 40), atypical EPEC (n = 29), ETEC (n = 27), EIEC (n = 14), typical EPEC (n = 5), EAEC/ETEC hybrids (n = 5), and EHEC (n = 2).

Pangenome characterization of our 248 EcoZUR pathogenic *E. coli* isolates identified a total of 23,516 orthologous genes (OGs). Of these, 3,021 OGs were present in more than 90% of the isolates, 11,060 OGs were found in less than 90% of the isolates, and 9,435 OGs were unique to a single isolate. We estimated the evolutionary relationships of the 248 isolates based on the sequences of OGs present in >99% of isolates (core Ogs) and built a core-genome maximum likelihood phylogeny (Figure 3). Initial phylogenomic characterization showed that all the 248 isolates circulating in Norther Ecuador were placed in six well-described phylogroups^62^ as follows: Phylogroups A (n = 104) and B1 (n = 62) harbored the most pathotypes, with B1 harboring all pathotypes and all of the EAEC/ETEC hybrids. Phylogroups B2 (n = 31) and D (n = 48) had a lower diversity of pathotypes and were largely dominated by isolates assigned to DAEC. Phylogroup C (n = 2) was the smallest phylogroup, housing only two DAEC isolates. A single isolate (B37-6) was placed in the cryptic phylogroup Clade IV, which we used as the outgroup.

**Figure 3:**
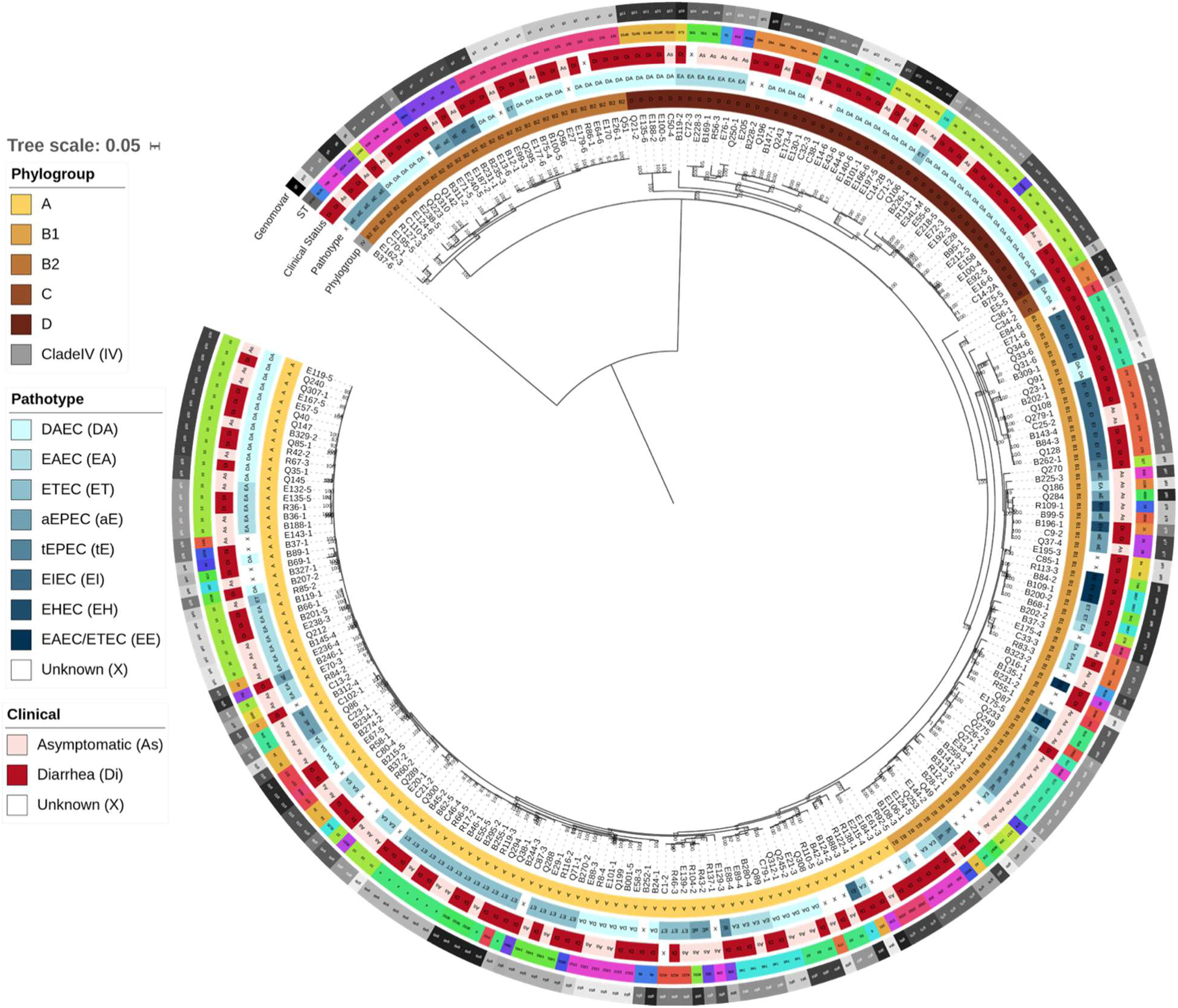
Distribution of EcoZUR pathogenic *E. coli* isolates: The core gene phylogeny is based on the SNP alignment of 376,389 base pairs from 2,577 core genes that were found in greater than 99% of the 248 high-quality genomes. The tree was constructed using RAxML (Stamatakis, 2014) v8.2.12 with the GTRGAMMA substitution model and was visualized using iTol (Letunic & Bork, 2021). The numbers displayed on the branches indicate the bootstrap values obtained after resampling the tree with replacement 1000 times. Phylogroup placement was first determined *in silico* using the tool ClermonTyping (Beghain et al., 2018) and further verified by their average nucleotide identity (ANI) values to members of the phylogroups. Phylogroup *in silico* assignments that did not fall into the majority phylogroup assigned based on ANI grouping were assumed unreliable and the phylogroup assignment was change to phylogroup in which it clustered best based on ANI. The pathotype for each genome was resolved by mapping high-quality reads to pathotype specific virulence genes (Table 1) and using greater than 1X sequencing depth to call the corresponding gene present (see Materials and Methods for more details).

We compared genomovar assignments to their placement on the core genome phylogeny and observed strong concordance between the genomovars defined by ANI clusters and the phylogenetic relationships inferred from core genes. Overall, 92% (46/50) of multi-genome genomovars were monophyletic in the core gene tree. The four exceptions—g34, g33, g38, and g49 all assigned to phylogroup A—exhibited partial paraphyly, with one member displaced from the primary cluster, likely reflecting minor inconsistencies between ANI-based clustering and the core gene phylogenetic signal (Figure 3). Nevertheless, the high concordance between genomovars and the core genome phylogeny further corroborates the robustness of our ANI-based clustering approach and the use of genomovars as biologically meaningful intra-species units.

The vast majority of genomovars were pathotype-uniform, with all member isolates sharing a single pathotype designation. However, notable exceptions revealed pathotype heterogeneity within individual genomovars (Figure 3). In genomovar g3, for example, a single isolate (Q295) was assigned to ETEC whereas the remaining members were classified as DAEC, with one genome unassignable by our diagnostic criteria. By contrast, the phylogenetically closest genomovar, g1—whose members were also designated ST-131 alongside those of g3—was mostly composed of DAEC, with a single genome unassignable to a pathotype, suggesting that the ETEC-associated virulence determinants for that single isolate were acquired after divergence from the g1 lineage. It is equally plausible, however, that the isolate lost its DAEC-associated determinants, with the ETEC designation reflecting a combination of gene gain and loss rather than a single acquisition event. Notably, pathotype discordance was not limited to single isolates within an otherwise uniform genomovar. In g94, for instance, three genomes sharing a mean ANI of 99.72% (± 0.016) received entirely discordant pathotype assignments: one as an EAEC/ETEC hybrid, one as aEPEC, and one that could not be classified under our established pathotype methods. That such divergent pathotype profiles can arise among genomovar members differing by approximately 0.3% ANI implies that the virulence determinants defining these pathotypes were most likely recently acquired (or lost) via horizontal gene transfer, with the unassignable isolate potentially representing an intermediate genomic state. We further corroborated the role of recent horizontal gene transfer based on phylogenetic analysis below.

### Genomovars expose rapid DEC pathotype switching driven by horizontal gene transfer

The above observations prompted us to ask whether the pathotype discordance detected within genomovars could be traced to discrete horizontal transfer events. To test this, we constructed tanglegrams to evaluate topological incongruences between the core genome phylogeny and individual gene trees generated from select pathotype-diagnostic markers (Table 1), providing a framework to identify signatures of lateral gene flow (Supplementary Figure 1). We centered our subsequent analysis on the DAEC pathotype—utilizing the *afa/dr* operon as the diagnostic locus—because it provided the most robust and compelling topological evidence for horizontal transfer. Genes associated with the *afa/dr* operon were arranged in a collinear configuration on a single contiguous sequence (contig) across 67 DAEC-assigned genomes. The tanglegram (Figure 4) revealed striking topological incongruence between the two trees: genomovars g1 and g3, both assigned to ST-131 and sharing a mean genome-wide ANI of 99.34% (± 0.07%), yet their *afa/dr* operon variants share only 98.40% nucleotide identity. Over the 4,305-bp operon sequence, this amounts to ∼69 substitutions, 2.4-fold more than the ∼29 substation expected if the operon were diverging under a similar rate as the genome average (i.e., the number of substitutions predicted by applying the genome-wide ANI to the operon length). Furthermore, the a*fa/dr* operon sequence for an isolate (B124-6) assigned to a different genomovar (g6) shared 100% sequence identity to all g3 members, yet its mean genome-wide ANI compared to g3 members was 98.35% (± 0.06%)—a level of genome divergence that would predict approximately 68 SNPs across the operon under similar expectations as above. In both g12 and g16 (intra-genomovar mean ANI: 99.79% ± 0.14 and 99.74% ± 0.14, respectively), a single isolate carried an *afa/dr* operon variant that was divergent from its genomovar members (operon sequence identity: 98.96% and 98.33%, respectively) yet more closely matched *afa/dr* sequences from phylogenetically distant genomovars (Figure 4).

**Figure 4.**
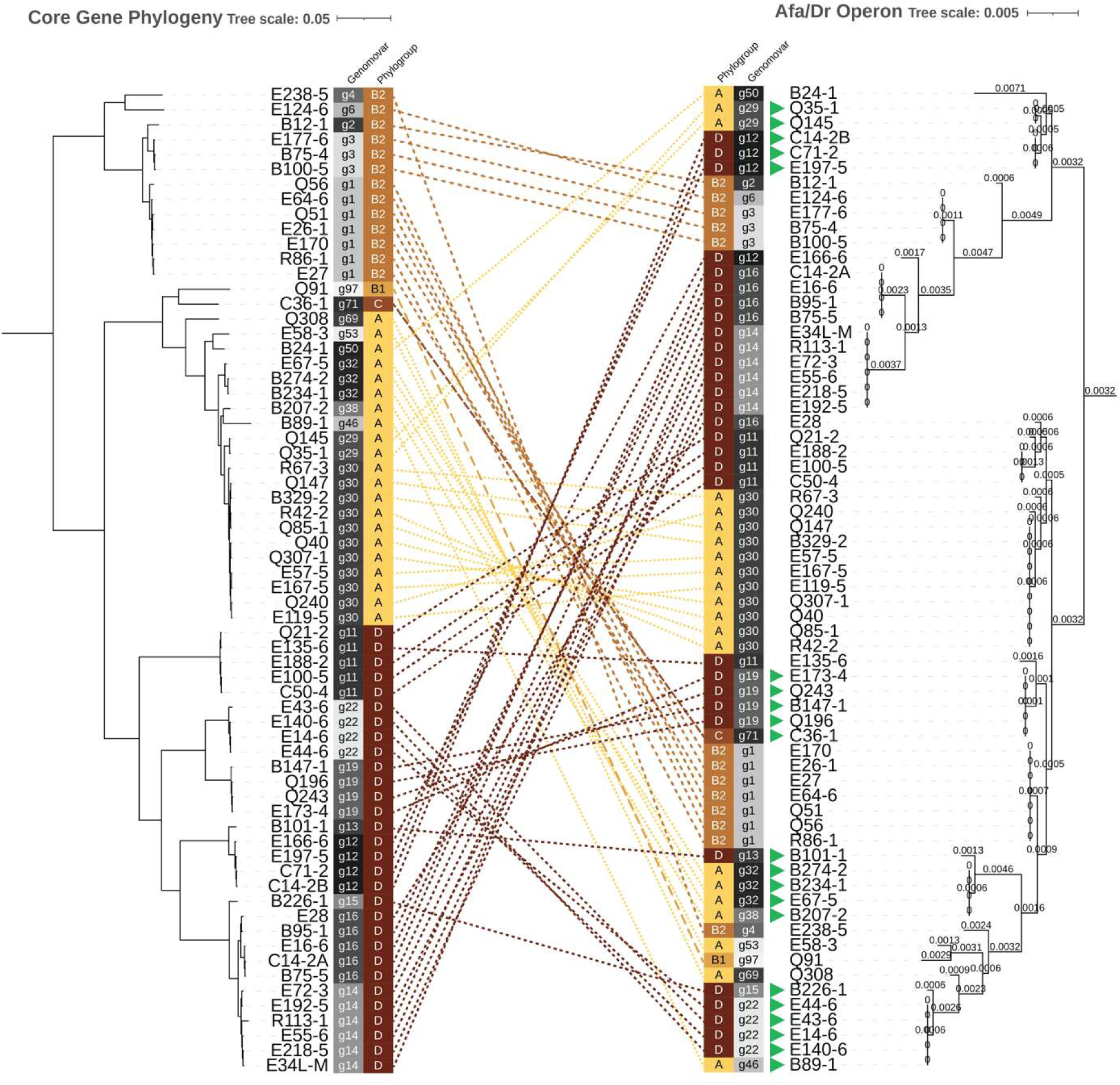
Phylogenetic incongruence identifies putative horizontal hene transfer in EcoZUR Diffuse Adherent *E. coli* (DAEC) isolates. Tanglegram comparing a pruned core gene tree (left) from Figure 1 with the Afa/Dr operon tree (right) of EcoZUR DAEC genomes, all of which contain complete genes on a single contig and exhibit collinear gene arrangements. Lines connecting corresponding taxa represent the same genome in both trees and are colored based on phylogroup. Both trees display the phylogroup and genomovar designations for each genome. The branch lengths in the operon tree represent the number of nucleotide substitutions that have occurred along each branch, as estimated using the Generalized Time Reversible (GTR) model with an approximate-maximum likelihood method. Green triangles in the operon tree represent clusters of genomes with identical or nearly identical sequences (i.e., fewer than five segregating sites across the operon) and include at least one genome from a different phylogroup.

We analyzed representative isolates from genomovars g19 and g71 (inter-genomovar mean genome ANI=97.07 ± 0.0311) by comparing the percent sequence identities of reciprocal best-matching genes (RBMs) centered around the *afa/dr* Operon (Figure 5). Although core and accessory RBMs exhibited sequence identities reflective of the substantial phylogenetic divergence between representative members from genomovars g19 and g71, the *afa/dr* operon was strikingly conserved, displaying 100% sequence identity across all representative genomes (Figure 5, top four panels). Notably, mobile genetic elements (MGEs) flanking the *afa/dr* operon also exhibited near-identical sequence conservation across those isolates from distinct genomovars and divergent phylogroups, further reinforcing the evidence for recent lateral transfer of this genomic region.

**Figure 5:**
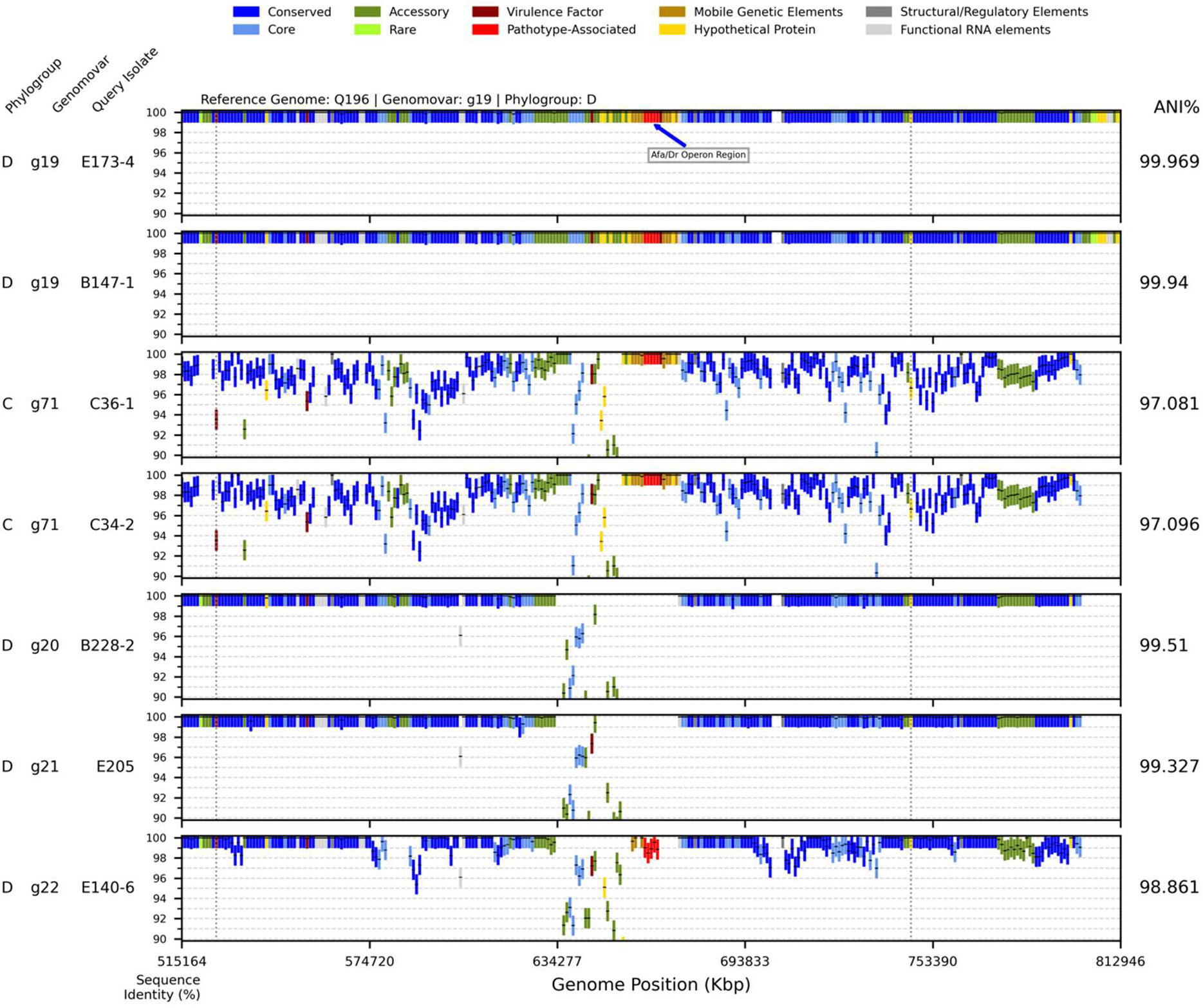
Horizontal gene transfer of the *afa/dr* operon between phylogenetically distant Diffuse Adherent *E. coli* (DAEC) isolates. Each panel displays reciprocal best-matching genes (RBMs) of a query genome mapped to reference genome Q196 (g19), centered on the *afa/dr* operon region and flanked by ∼150 genes upstream and downstream. RBMs are arranged by their relative genomic position on the reference (x-axis) with percent sequence identity on the y-axis; grey vertical dotted lines denote contig boundaries. Phylogroup, genomovar, and genome ANI to the reference are indicated for each query. RBMs are color-coded by functional category (see Methods). Panels 1–2: Genomes from the same genomovar as the reference (E173-4 and B147-1, g19) show uniformly high (∼100%) sequence identity across all RBMs, reflecting within-genomovar conservation. Panels 3–4: Genomes from a different phylogroup and genomovar (C36-1 and C34-1, g71) show expected divergence across conserved, core, and accessory genes, yet the *afa/dr* operon and flanking MGEs retain identical or near-identical sequences—strong evidence of recent HGT between distantly related lineages. Panels 5–6: Genomes from the same monophyletic clade as the reference but assigned to different genomovars (B228-2, g20; E205, g21) exhibit high identity across most RBMs but completely lack the *afa/dr* operon and associated MGEs, suggesting lineage-specific loss or absence of horizontal acquisition. Panel 7: Genome E140-6 (g22), from the same phylogroup as the reference, carries the *afa/dr* operon but with greater sequence divergence and absence of several flanking MGEs, consistent with an independent HGT acquisition from a different donor. Together, these comparisons demonstrate that the *afa/dr* operon has been horizontally transferred on multiple independent occasions across phylogenetically diverse *E. coli* lineages.

Synteny analysis of representative genomes from both closely and distantly related genomovars further corroborated this pattern (Supplementary Figure 2). Distantly related genomovars g19 and g71 displayed remarkably conserved synteny of the MGEs flanking the *afa/dr* operon, consistent with the recent acquisition of a shared mobile cassette. In contrast, closely related genomovars—both at the inter-genomovar (g1 vs. g2 vs. g3) and intra-genomovar level (g16 and g12)—exhibited variable MGE composition and arrangement flanking the operon, suggesting independent insertion events and ongoing genomic rearrangement (Supplementary Figure 2). Across all isolate genomes analyzed in Figure 4, the *afa/dr* operon was consistently embedded within a dense and diverse repertoire of MGEs on both sides, including transposase-encoding sequences (*insI1, istA, istB, tnpA, tnpB*, and *tra5*), members of numerous insertion sequence (IS) families (IS*1*, IS*2*, IS*3*, IS*4*, IS*21*, IS*30*, IS*66*, IS*629*, IS*As1*, IS*Cro1*, IS*Ec10*, IS*Ec52*, IS*L3*, IS*110*, IS*200*/IS*605*, and IS*682*), integrase-encoding sequences—most notably the class 1 integron integrase *intI1*—and several uncharacterized hypothetical proteins. Genomic origin prediction across all isolates analyzed in Figure 4 revealed that approximately 95% of the contigs harboring the *afa/dr* operon were of chromosomal origin, with the remaining 5% predicted to be plasmid-borne (data not shown). Taken together, these findings are consistent with previous reports demonstrating that the *afa/dr* operon is characteristically embedded within a conserved MGE-rich genomic neighborhood and can reside on either chromosomal genomic islands or plasmids^63–65^ providing strong support for the capacity of this operon to mobilize and disseminate via lateral gene transfer. However, the extent to which these horizontal gene transfers directly influence pathogenic potential remains unclear, as additional factors such as regulatory changes, genetic background, and selective pressures likely modulate their functional impact.

### Genomovar-level virulence profiles reveal hierarchical phylogroup structure and hints at lineage-dependent pathogenic potential in diarrheagenic E. coli

We identified 701 virulence factors (VFs) across 247 *E. coli* genomes and classified them into ten functional categories: effector delivery systems (42.51%), adherence (26.11%), nutritional/metabolic factors (7.7%), motility (7.13%), immune modulation (6.28%), exotoxins (3.85%), invasion (2.99%), biofilm (0.86%), regulation (0.86%), exoenzymes (0.57%), and other (1.14%) (Supplementary Table 2). Based on their prevalence across the isolate genomes used in this study (Supplementary Figure 4A), VFs partitioned into four pangenome classes: core (≥90% of isolates; n = 74), accessory (>10–<90%; n = 171), rare (≤10%; n = 195), and singleton (n = 67) (Supplementary Figure 4B, Supplementary Table 2). Core VFs were enriched for functions essential to colonization and survival—motility, nutrient acquisition, and adherence—whereas accessory, rare, and singleton VFs spanned all functional categories, reflecting the substantial genetic plasticity of the *E. coli* accessory genome (Supplementary Figure 4C). Furthermore, we categorized 194 virulence factors as pathotype-specific (Supplementary Figure 4B), which encompassed the pathotype-diagnostic genes (Table 1) and additional well studied pathotype virulence determinants documented in the literature. Representative examples of those additional pathotype-specific virulence factors are discussed below.

EIEC genomes assigned to genomovar g99 and the singleton B88-3 (g66) carried multiple virulence genes associated with the invasion plasmid^1,66,67^ (pINV). These included effector delivery system genes belonging to the *ipa-mxi-spa* island and the *virB* gene, all of which are located within the conserved 31Kbp pINV entry region^8^. In contrast, five isolates assigned to EIEC by our read-mapping approach and classified within genomovar g100 lacked pINV-associated genes (Supplementary Figure 4D). In these isolates, the *ipaH* marker used for assigning pathotypes was predicted as chromosomally encoded rather than plasmid borne (data not shown), suggesting that reliance on a single *ipaH* gene may be insufficient for robust EIEC identification. Isolates assigned to aEPEC, tEPEC and EHEC pathotypes showed evidence of several virulence factors located on the locus of enterocyte effacement (LEE) pathogenicity island, including effector genes for the *esp* operon^68,69^. Of the five isolates identified as tEPEC, only three isolates assigned to the same genomovar possessed VFs for the complete *bfp* operon located on the *E. coli* adherence factor (EAF) plasmid^70^. Only a few EAEC *E. coli* pathotypes harbored VFs associated with the 55- to 65-MDa “pAA” plasmid^71^, genes that are critical for encoding aggregative adherence fimbriae (AAF) that facilitate biofilm formation, while the majority harbored genes belonging to the *aai* operon. In addition to the heat-labile enterotoxin^72^ (LT) *eltA* and *eltB* genes used to assign the pathotype, most ETEC isolates carried the plasmid-encoded type IV pilus called “Longus”, containing the majority of *lng* genes involved in pilus formation and adherence^73,74^, with several isolates without a pathotype assignment carrying *lng* genes. Although all DAEC assigned isolates harbored VFs related to the a*fa/dr* operon, only one isolate (B234-1) additionally carried the EAEC-associated *aai* operon (Supplementary Figure 3 and Supplementary Figure 4D), consistent with a potential DAEC/EAEC hybrid pathotype and highlighting the role of horizontal gene transfer in shaping the mosaic distribution of virulence determinants among co-circulating *E. coli* lineages.

To evaluate how virulence factors may affect the disease potential of genomovars, we restricted the current analysis to genomovars with at least three genome members, which included 33 genomovars comprised of a total of 162 isolate genomes across four known phylogroups. These criteria allowed us to assess how effectively genomovars captured shared virulence gene repertoires while minimizing the influence of random noise from singletons and genomovars with two members only.

We first assessed the relationship between genomic relatedness and accessory virulence factor composition among genomovars. A Mantel test comparing pairwise genome ANI distances against the binary (presence/absence) dissimilarity matrix of accessory virulence factors revealed a moderate positive correlation (r = 0.394, p < 0.001, n = 162), indicating that isolates sharing greater genomic similarity (i.e., being members of the same genomovar) also tended to harbor more similar virulence gene repertoires. Next, we tested whether phylogroup membership captured significant variation in accessory virulence factor composition and found that virulence gene repertoires were significantly differentiated by phylogroup (PERMANOVA: pseudo-F = 46.56, P = 0.001). Pairwise PERMANOVA comparisons between paired phylogroups further demonstrated that all pairs were significantly differentiated after correction for multiple testing (p-adj<0.006 for all comparisons), with the largest effect sizes observed between phylogroups B2 and D (pseudo-F = 121.13) and between B1 and D (pseudo-F = 80.56). However, significant heterogeneity in multivariate dispersion among phylogroups (PERMDISP: F = 20.99, P = 0.001) indicated that a portion of the overall PERMANOVA signal may reflect unequal within-group variance rather than differences in group centroids alone. Pairwise PERMDISP tests revealed that this heterogeneity was not uniformly distributed: genomes forming phylogroups A and B1 did not differ significantly in dispersion (p-adj = 0.449), nor did B2 and D (p-adj = 0.449), whereas all other phylogroup pairwise comparisons were significant (p-adj < 0.006). In other words, phylogroups B2 and D shared a comparably narrow range of accessory virulence gene repertoires across their constituent genomovars, whereas phylogroups A and B1 were characterized by broader, more variable virulence gene content, indicating two distinct modes of virulence gene organization among EcoZUR *E. coli* lineages. This structure was readily apparent in the UMAP reduced space of isolate-level virulence profiles, in which isolates segregated primarily by phylogroup and further resolved into discrete genomovar clusters, indicating that accessory virulence gene content is largely conserved at the genomovar level and structured according to phylogroup affiliation (Figure 6). Accordingly, genomes assigned to phylogroups B2 and D formed comparatively tight, well-resolved clusters, whereas those assigned to phylogroups A and B1 exhibited substantially greater dispersion, suggesting that the latter harbor more heterogeneous accessory virulence gene repertoires across their constituent genomovars (Figure 6). This interpretation was further supported by observing the relative distribution of virulence factor functional categories across genomovars (Supplementary Figure 5).

**Figure 6:**
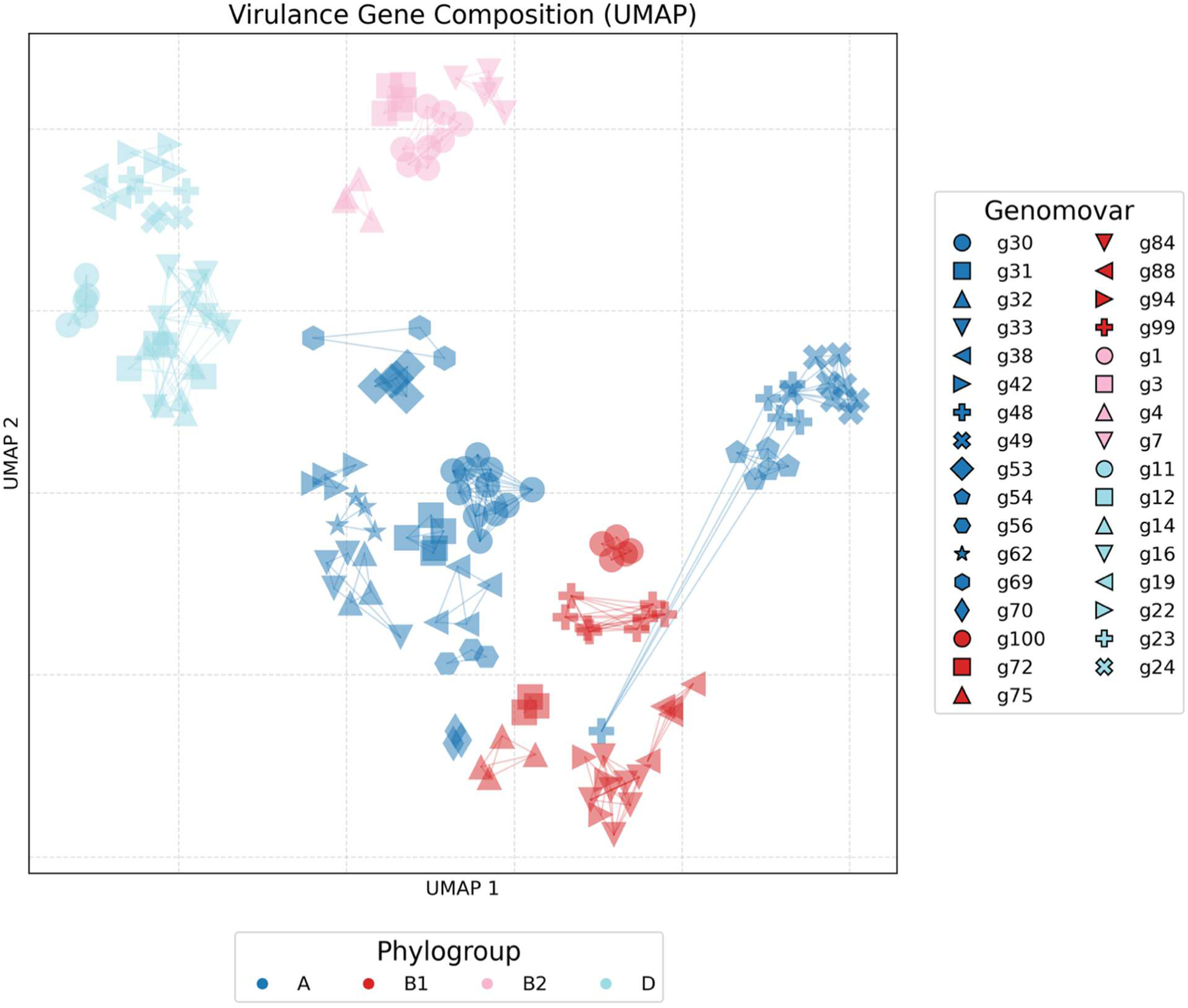
Uniform Manifold Approximation and Projection (UMAP) visualization of accessory virulence factor repertoires reveals phylogroup-specific clustering with genomovar substructure. Two-dimensional projection of pairwise Dice-Sørensen dissimilarities calculated from accessory virulence factor (VF) presence-absence profiles across genomovar clusters containing three or more genomes (n = 162). Each point represents an individual isolate positioned in UMAP space with color indicating phylogroup assignment and shape denoting genomovar cluster membership. Isolates exhibit separation by phylogroup, reflecting significant global differentiation in accessory VF repertoires (PERMANOVA: pseudo-F = 46.56, p = 0.001), with finer-scale clustering patterns corresponding to genomovar assignments within phylogroups. This nested structure demonstrates that while accessory virulence gene architecture is primarily shaped by phylogenetic background, additional strain-level variation contributes to within-phylogroup diversification of virulence potential.

Notably, isolates assigned to phylogroups B2 and D—whose virulence profiles were the most tightly conserved—also exhibited the highest proportions of diarrhea-associated isolates compared with phylogroup A (Supplementary Figure 6). This pattern was particularly striking among DAEC isolates, the most abundant pathotype in our EcoZUR collection and conventionally regarded as less virulent^75,76^. DAEC genomes assigned to phylogroups B2 and D showed markedly higher disease-associated proportions (82.1% and 74.2%, respectively) than those assigned to phylogroup A (52.1%), despite sharing the same pathotype designation (Supplementary Figure 5). The odds of disease association were approximately two-fold higher for both B2 and D relative to phylogroup A (OR = 1.96 and 1.85, respectively), though these differences did not reach statistical significance (Fisher’s exact test: p = 0.307 and p = 0.207, respectively).

We next asked whether rare gene content could account for differences in disease association between closely related genomovars. To this end, we selected pairs of genomovars sharing high genomic relatedness but exhibiting contrasting proportions of diarrhea-associated or asymptomatic (control) isolates and examined whether specific rare virulence genes were differentially distributed between them. To minimize confounding from pathotype-associated gene content, we restricted comparisons to genomovar pairs sharing the same pathotype designation and evaluated the union of their rare virulence genes, identifying those exclusive to one genomovar versus the other. We then extended this analysis across all genomovar clusters by constructing a presence/absence heatmap of the identified rare genes to assess whether their distribution corresponded with differences in pathogenic potential.

Supplementary Figure 7 illustrates this approach, comparing genomovar g31 (lower disease-associated genomes) and g42 (higher disease-associated isolates), both belonging to the EAEC pathotype and assigned to ST-10 and sharing a mean genome-wide ANI of 99.23% (SD = 0.051). Notably, most g42 members carry a suite of predicted virulence genes absent in g31. For example, g42 genomes contain genes encoding putative effector delivery system components (EC55989 prefix) alongside multiple type VI secretion system (T6SS) genes (*tssA*, *tssE*, *tssH*, and *tssK*). Because the T6SS delivers effectors that destabilize host cells and intoxicate competing bacteria^77^, the presence of these genes may enhance T6SS-mediated niche competition and host interactions, potentially driving the increased virulence potential of g42 isolates. To assess whether the pattern of these rare genes and disease association extended beyond the g31–g42 comparison, we examined the distribution of these genes across additional genomovar clusters. Several clusters that broadly harbored these genes (including g62, g70, and g75) showed similarly elevated proportions of diarrhea-associated isolates. In contrast, genomovar clusters g31 and g24 largely lacked these genes and exhibited markedly lower proportions of diarrhea-associated members.

## Discussion

The conventional classification of diarrheagenic *E. coli* (DEC) into discrete pathotypes^1^ has served as a foundational framework for clinical microbiology and epidemiological surveillance. Yet this scheme implicitly treats pathotype identity as a stable lineage property—an assumption increasingly difficult to reconcile with the genomic fluidity revealed by population-scale sequencing. By applying the recently described intra-species ANI gap^17^ approach to resolve 248 *E. coli* isolates from the EcoZUR cohort into genomovars, we demonstrate that the natural genomic discontinuity at approximately 99.5% ANI delineates intra-species units that are both phylogenetically coherent and functionally/epidemiologically informative. These units reveal cryptic population structure that conventional MLST cannot resolve. In our dataset, 14 STs (including ST-10 and ST-131) each encompassed multiple genomovars, indicating that a single sequence type can harbor substantial intra-ST divergence. Conversely, 14 genomovars each contained genomes assigned to more than one ST, showing that distinct ST designations can collapse into a single genomovar. This discordance is not merely a technical artifact of marker gene resolution; it reflects the fundamental difference between classification systems anchored to a handful of slowly evolving (relatively to the genome average) housekeeping loci and one that integrates signal from thousands of shared genes and a natural discontinuity in genomic diversity (as opposed to an arbitrary sequence identity threshold). The observation that this intra-species discontinuity (∼99.5% ANI) has now been documented across hundreds of bacterial species and even in bacteriophage and eukaryotic viruses^17,78^ suggests that the genomovar boundary captures a universal feature of microbial population biology. Recent work has provided a mechanistic basis for this boundary, demonstrating that members of the same genomovar—which share high sequence identity and tend to co-occupy the same ecological niche—engage in frequent, genome-wide homologous recombination that purges accumulated mutations and actively maintains genomic cohesion and phenotype within the genomovar unit^79^. Because homologous recombination efficiency scales with sequence identity, this cohesive force operates most effectively among the closely related genomes within a genomovar. Between more divergent genomovars, recombination becomes less efficient and progressively acts instead as a barrier to species (or population, more generally) convergence as sequence identity decreases below the 95% level, which introduces heterologous segments and drives rapid divergence from the parent species/population^79^. The ANI gap (at approximately the 99.5%) observed here among the EcoZUR genome is highly consistent with these previous results and interpretations. Overall, our findings underscore how genomovar delineation can identify biologically natural populations better positioned to capture differences in genome content, including horizontally acquired virulence determinants, antimicrobial resistance genes, and pathogenicity islands. This framework therefore offers a more reliable basis than MLST alone for distinguishing functionally relevant differences among closely related lineages, even when those lineages share core genome architecture.

The most striking biological insight from our genomovar-resolution analysis is the rapidity with which pathotype identity can change. Pathotype discordance within individual genomovars indicates that pathotype identity can change over remarkably short evolutionary timescales. For example, the three members of genomovar g94 differ by only ∼0.3% ANI yet represent three distinct pathotypes, implying that the horizontally acquired loci defining pathotype identity can be gained or lost on timescales far shorter than those required for appreciable core genome divergence or those captured by a shared sequence type designation. Because such transitions occur within the diversity of a single genomovar and within a narrow sampling window (notably, all isolates in this study were recovered over an approximately 18-month period across northern Ecuador^18^), they underscore the role of lateral gene acquisition in driving rapid, ongoing shifts in pathotype (and virulence) potential among co-circulating *E. coli* populations. Furthermore, the tanglegrams analysis (Figure 4 and Supplementary Figure 1), synteny analysis of the *afa/dr* operon (Supplementary Figure 2), and the locus-centered identity plots (Figure 5 and Supplementary Figure 3) provide a mechanistic basis for this plasticity. For instance, the *afa/dr* operon’s consistent embedding within a dense, heterogeneous repertoire of mobile genetic elements—including IS families, transposases, and the class 1 integron integrase *intI1*—effectively transforms it into a modular virulence cassette capable of chromosomal integration across phylogenetically disparate lineages. The finding that phylogenetically distant genomovars from different phylogroups (e.g., g19 and g71) can share identical *afa/dr* and flanking MGE sequences, while closely related sister genomovars (e.g., g1 and g3) carry divergent operon variants with substitution rates exceeding expectations from average core genome divergence by more than two-fold, argues that the evolutionary dynamics of this locus are dominated by lateral acquisition and replacement rather than vertical inheritance and gradual divergence (or point mutation). Furthermore, a single DAEC-assigned genome (B234-1) harbored a near complete EAEC-associated AAI operon (that was absent from the remaining members), generating a DAEC–EAEC hybrid and illustrating that HGT-driven plasticity can encompass entire virulence modules (Supplementary Figure 3). Such bulk transfer of virulence cassettes highlights a profound mechanism by which discrete horizontal transfer events rapidly diversify the pathogenic potential of individual strains. If pathotype-defining loci behave as mobile accessories rather than stable genomic fixtures, then pathotype designations capture the instantaneous virulence configuration of a genome rather than a long-lasting evolutionary identity—a distinction with profound implications for how we interpret the epidemiology of DEC. In endemic settings such as northern Ecuador, where extensive spatial mixing has already been documented^16^, this lateral mobility may enable the rapid, recurrent emergence of novel pathotype configurations from a shared pool of co-circulating lineages, potentially explaining why disease associations have often proven so difficult to attribute to individual clones or pathotypes. Overall, these findings suggest that horizontal gene transfer may facilitate shifts in pathotype designation by enabling the rapid acquisition of virulence-associated genes, potentially accelerating the transition from commensal to pathogenic lifestyles or allowing *E. coli* to switch between pathogenic pathotypes relatively quickly.

Our analyses further reveal that the phylogenetic background of a lineage—independent of pathotype designation—appears to modulate virulence potential in ways that the pathotype framework alone cannot capture. Accessory virulence gene repertoires were significantly structured by phylogroup, with genomovars assigned to phylogroups B2 and D exhibiting tightly conserved, functionally uniform virulence profiles, whereas those in phylogroups A and B1 harbored substantially more heterogeneous accessory gene content across their constituent genomovars (Figure 6). This dichotomy may reflect fundamentally different evolutionary strategies: genomovars in phylogroups B2 and D, which are enriched for extra-intestinal pathogenic *E. coli* (ExPEC) lineages and have been associated with urinary tract infections, bacteremia, and neonatal meningitis^1,80–82^, may maintain a more constrained virulence architecture due to strong(er) purifying selection in the context of systemic infection, where a precise repertoire of immune evasion, iron acquisition, and adhesion factors is required for host colonization outside the gut. In contrast, the broader accessory virulence gene diversity observed across genomovars within phylogroups A and B1 likely reflects, at least in part, the substantially greater pathotype diversity harbored by these phylogroups—B1 alone encompassed all pathotypes detected in our collection, including the EAEC/ETEC hybrids—indicating that a wider array of pathotype-specific virulence loci contributes to the accessory gene pool and inflates inter-genomovar dissimilarity. This interpretation is consistent with the well-documented role of phylogroups A and B1 as predominantly commensal lineages that exhibit high genomic diversity and broad ecological distribution across human and animal hosts^83,84^. The observation that DAEC isolates assigned to phylogroups B2 and D exhibited markedly higher disease-association rates (82.1% and 74.2%) than those in phylogroup A (52.1%), despite sharing an identical pathotype designation, lends empirical support to the hypothesis that phylogroup-specific differences in accessory virulence gene content modulate pathogenic potential even within a single pathotype. Such lineage-associated variation in virulence capacity may contribute substantially to the inconsistent clinical outcomes historically reported for DAEC and warrants further investigation with larger, epidemiologically matched cohorts to determine whether the trend observed here achieves statistical significance at greater sample sizes.

The identification of rare virulence gene signatures—such as those putative effector delivery system genes belonging to the EC55989 family—that distinguished disease-associated from asymptomatic genomovars within the same pathotype is intriguing and raises the possibility that specific rare gene acquisitions may tip the balance from commensal carriage to overt disease. However, we caution that interpretation of these rare gene differences proved challenging for several reasons. Many of the detected genes correspond to predicted entries in the VFDB that have not been functionally characterized experimentally, and their designation as virulence factors remains provisional. While these genes were predominantly observed in genomovars assigned to the EAEC pathotype (Supplementary Figure 7), this association may reflect artifacts of limited sample sizes within pathotypes rather than true biological specificity. We cannot exclude the possibility that the combinatorial effect of multiple virulence genes—including the broader accessory virulence repertoire and pathotype-associated loci—rather than the presence of individual rare genes, underlies differential disease potential. Moreover, our analysis was based solely on gene presence–absence patterns of virulence genes and does not account for allelic diversity within these loci, which may also modulate virulence phenotypes through altered expression, protein function, or regulatory context. These limitations complicate efforts to attribute pathogenic capacity to specific loci based on presence–absence patterns alone, and thus to definitively classify genomovars as virulent versus avirulent. Taken together, while these findings raise the intriguing possibility that rare gene content differences may harbor genuine virulence determinants, they also underscore the inherent difficulty in disentangling lineage-specific, pathotype-associated, and virulence-linked genetic variation using comparative gene content approaches alone.

In conclusion, our study provides the first genomovar-resolution portrait of diarrheagenic *E. coli* population dynamics within an endemic setting and reveals three properties of *E. coli* biology that the conventional pathotype framework obscures. First, pathotype identity is labile: it can be rewritten by a single horizontal transfer event on timescales shorter than those required for measurable core genome divergence, rendering pathotype designations informative snapshots of a genome’s current virulence configuration rather than stable evolutionary markers. Second, the phylogenetic chassis matters: the genomic background in which horizontally acquired virulence loci are received shapes their phenotypic expression, such that two isolates sharing the same pathotype but belonging to different phylogroups may present fundamentally different disease risks. Third, the genomovar framework offers a natural, reproducible unit for comparative pathogenomics that complements MLST in resolving functionally relevant genomic diversity and provides a scalable foundation for linking intra-species genomic variation to clinical and epidemiological outcomes. Future work integrating transcriptomic and functional data with genomovar-level classification will be essential to determine which of the virulence gene signatures identified here translate into measurable phenotypic differences—and ultimately, to develop genomics-informed risk stratification tools for diarrheagenic *E. coli* surveillance in endemic regions.

## Acknowledgments

We thank all participants who provided data and samples for the EcoZUR study, the Ecuadorian Ministry of Public Health for providing access to their clinics and hospitals, and the local field and laboratory teams for assistance in collecting and processing samples. Funding for this study was provided by the National Institute of Allergy and Infectious Diseases grants 1K01AI103544 (to K.L.), by the National Institute for Environmental Health Sciences (5T32ES007032 and 5T32ES102870 to K.J.J.), and by the National Science Foundation award 2129823 (to K.T.K.). Raw reads were deposited in NCBI Sequence Read Archive (BioProject ID PRJNA486009). The content of this paper is solely the responsibility of the authors and does not necessarily represent the official views of the National Institutes of Health or the National Science Foundation.

## Supplementary material

**Supplementary Figure 1:**
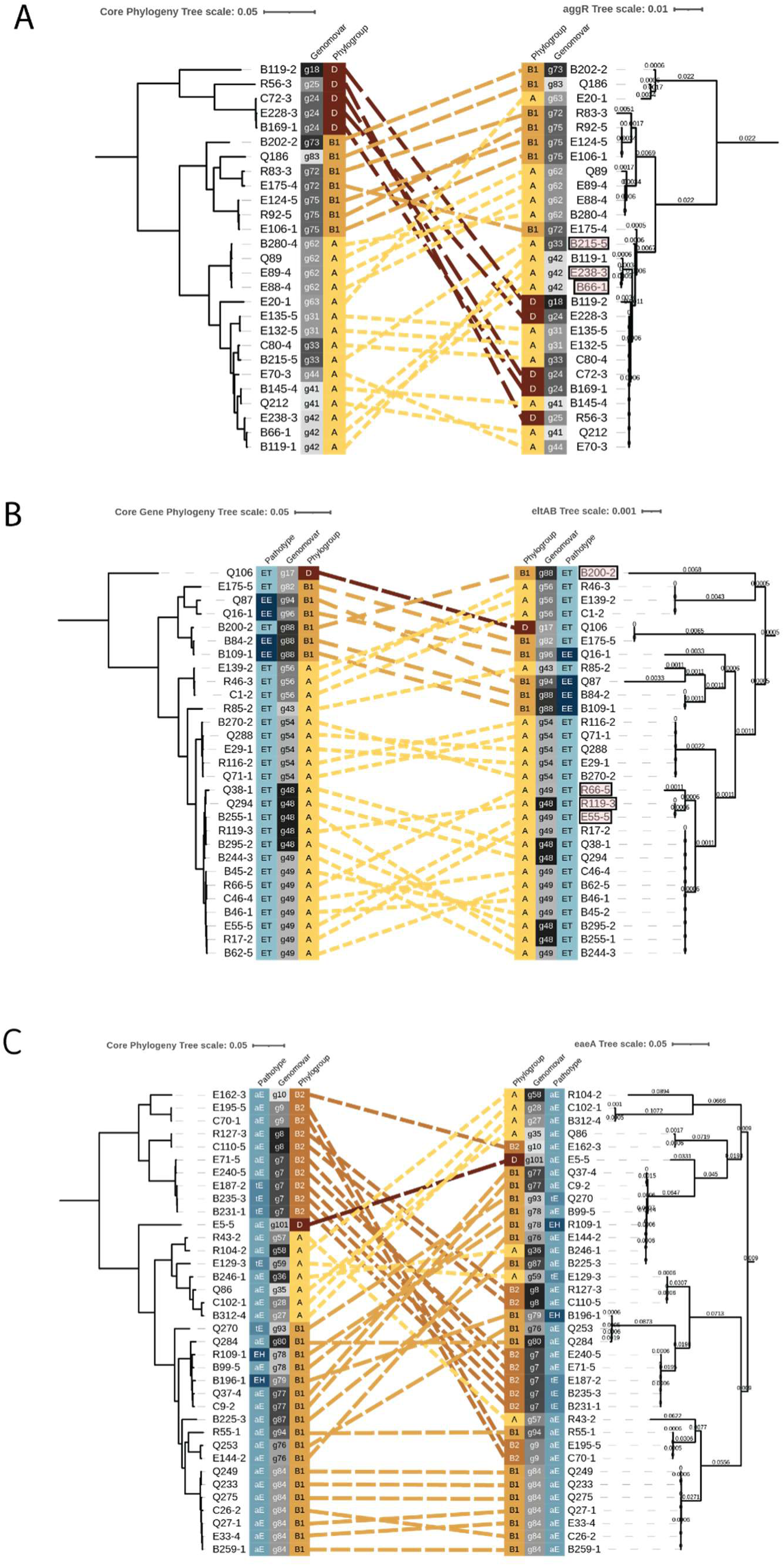
Phylogenetic incongruence reveals horizontal transfer of pathotype-diagnostic genes. Tanglegrams comparing the core genome phylogeny (left) with approximate-maximum likelihood trees of specific diagnostic virulence loci (right) from Table 1. Lines connect identical isolates across the two phylogenies. (A) Tanglegram for aggR gene used to define Enteroaggregative *E. coli* (EAEC) pathotype. (B) Tanglegram for eltAB genes (concatinated) used for defining Enterotoxigenic E. coli (ETEC). (C) Tanglegram eaeA gene, a locus shared among atypical Enteropathogenic (aEPEC), typical Enteropathogenic (tEPEC), and Enterohemorrhagic E. coli (EHEC) pathotypes. The branch lengths in the gene trees represent the number of nucleotide substitutions that have occurred along each branch, as estimated using the Generalized Time Reversible (GTR) model with an approximate-maximum likelihood method. Genomes highlighted with a red box indicate an incomplete diagnostic gene sequences relative to the reference.

**Supplemantary Figure 2:**
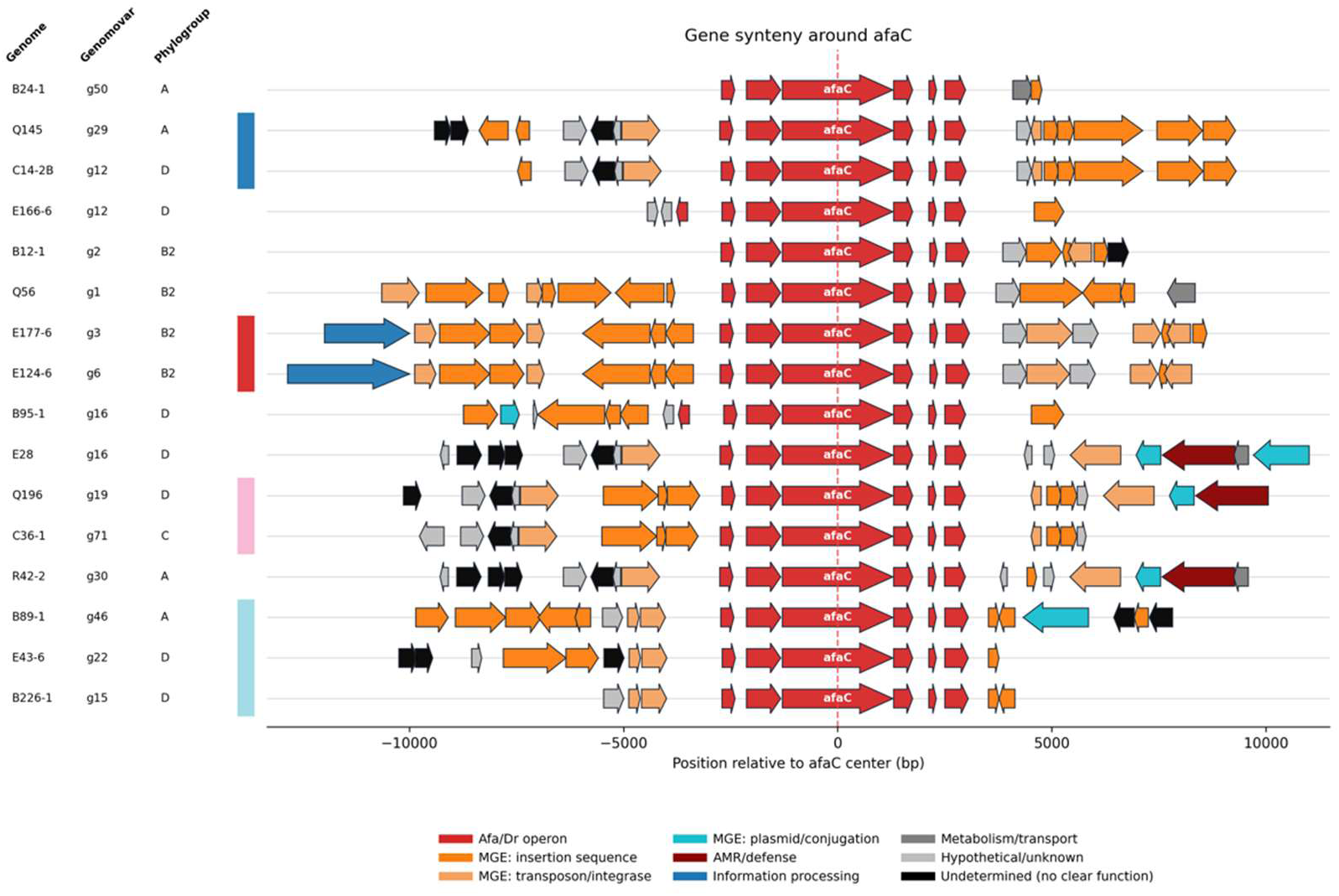
Gene synteny around the afaC (target) locus was visualized by parsing Bakta (v1.9) genome annotation outputs, extracting coding sequences within a defined flanking window (±10 genes) of the target gene, and converting genomic coordinates to positions relative to the target gene center. Each gene was classified into a functional category based on gene name and product annotation, and rendered as a directional arrow coloured by category using matplotlib (v3.x) in Python 3.12.12. Vertical colored bars represent distently related genomes (ANI <98.5%) from distncit genomovars sharing identical (or nearly idential) *afa/dr* orperon and mobile genetic element (MGE) sequences, which are also represented in Figure 4 (green triangles).

**Supplementary Figure 3:**
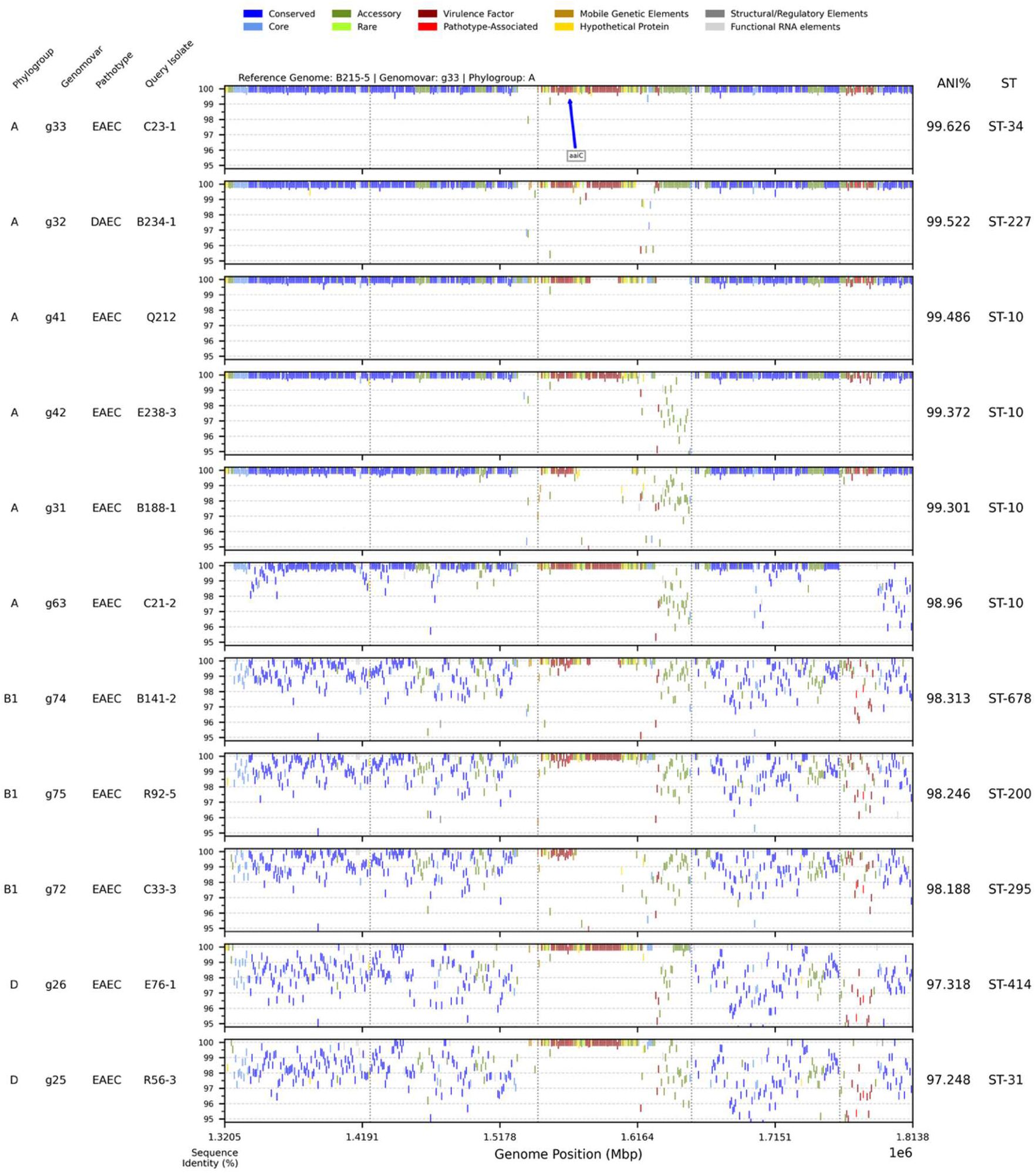
Sequence identity and structural conservation of the laterally acquired aaiC-associated operon in isolate B234-1. Comparative locus-centered projection detailing the genomic architecture of the full virulence cassette surrounding the EAEC-diagnostic *aaiC* gene. Predicted coding sequences were compared to a reference contig (∼115-kbp) harboring *aaiC* using reciprocal best-match alignments. The plot displays the reference genomic position (x-axis) against the nucleotide sequence identity of aligning orthologs (y-axis). Gene models are colored by functional and pangenome categories (refere to methods) to highlight the physical co-localization of virulence markers with mobile genetic elements (MGEs) and hypothetical proteins.

**Supplementary Figure 4.**
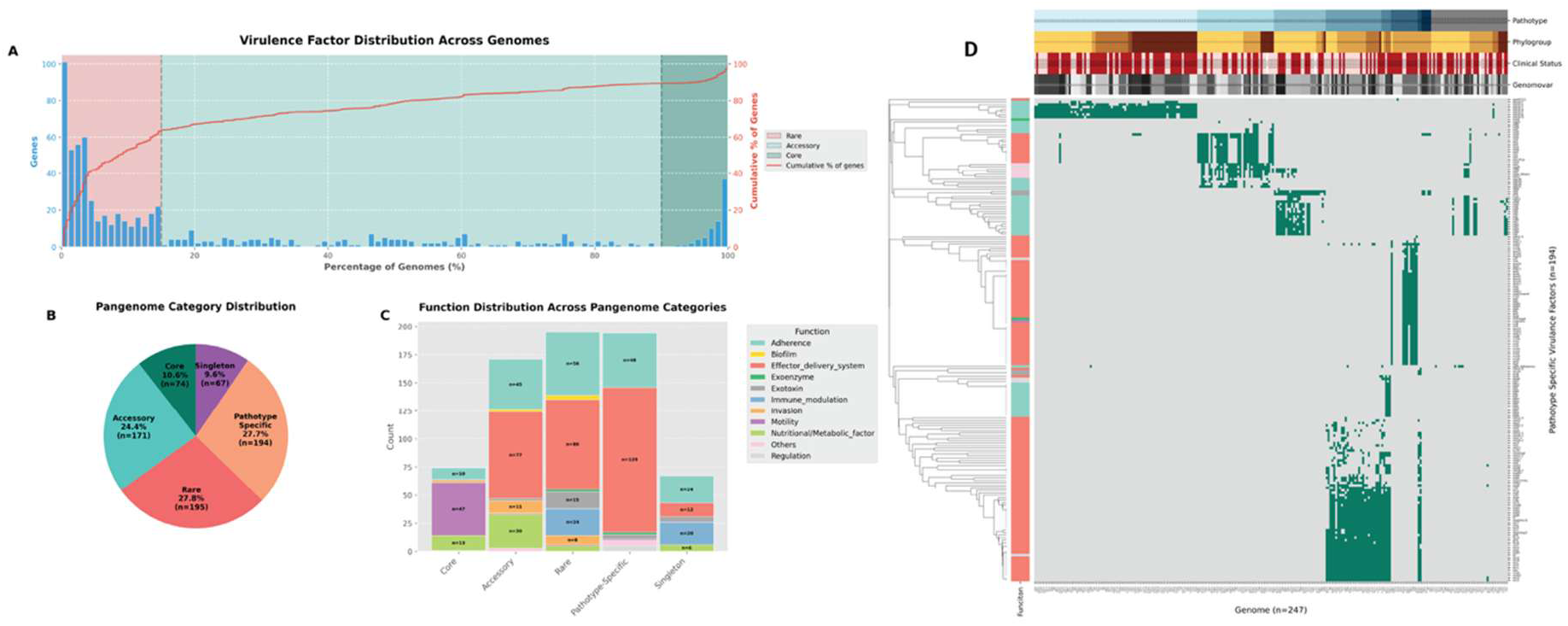
Distribution and functional composition of virulence factor genes across the *E. coli* EcoZUR genomes. **(A)** Cumulative distribution of virulence factors (VFs) showing the relationship between VF prevalence and the percentage of genomes harboring genes. VFs were classified as rare (≤15%, but not including singletons), accessory (15-90%), or core (≥90%) based on their frequency distribution across the population. The overlaid curve shows the cumulative frequency. (**B)** Proportional distribution of VFs across pangenome categories. VFs were classified as pathotype-specific (such as diagnostic genes defined in Table 1 and genes shown to be association with a given pathotype by reviewing peer-reviewed literature) or by virulence pangenome category (rare, accessory, core, singleton). Values represent the percentages (and counts) of VFs in each classification. **(C)** Functional composition of VFs within each pangenome category. Bars represent the total count of VFs pangenome classification as core with colors indicating functional categories as annotated in the Virulence Factor Database (VFDB), illustrating how the biological roles of virulence determinants vary with their genomic pangenome distribution classification. **D).** Cluster map illustrating the presence (green) of pathotype-specific virulence factors among genomes exhibiting >95% ANI. Annotations for functional category, pathotype, phylogroup, clinical status, and genomovar are indicated to contextualize the gene distribution.

**Supplementary Figure 5:**
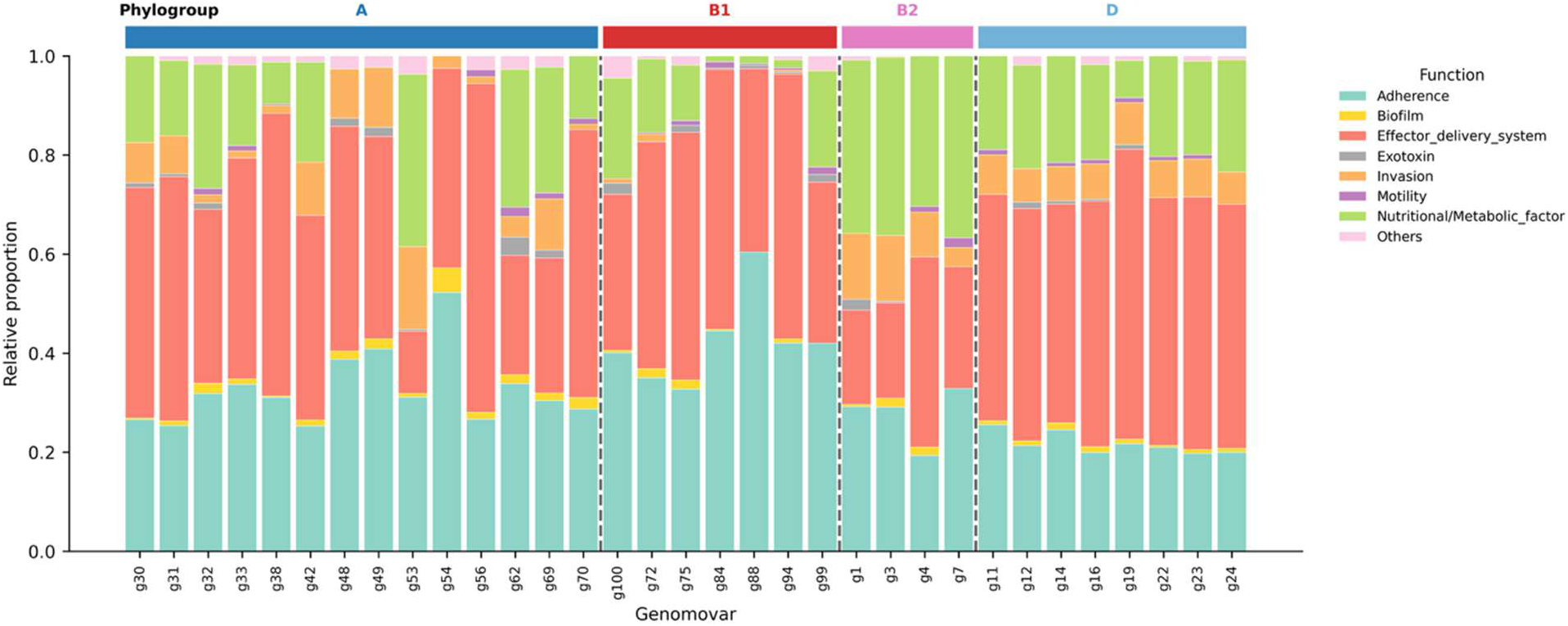
Virulence factor functional composition across genomovars and phylogroups. Stacked bar plots showing the relative proportions of virulence factor functional categories for each genomovar (see figure key), grouped by phylogroup (designated by the corresponding letter on the top).

**Supplementary Figure 6:**
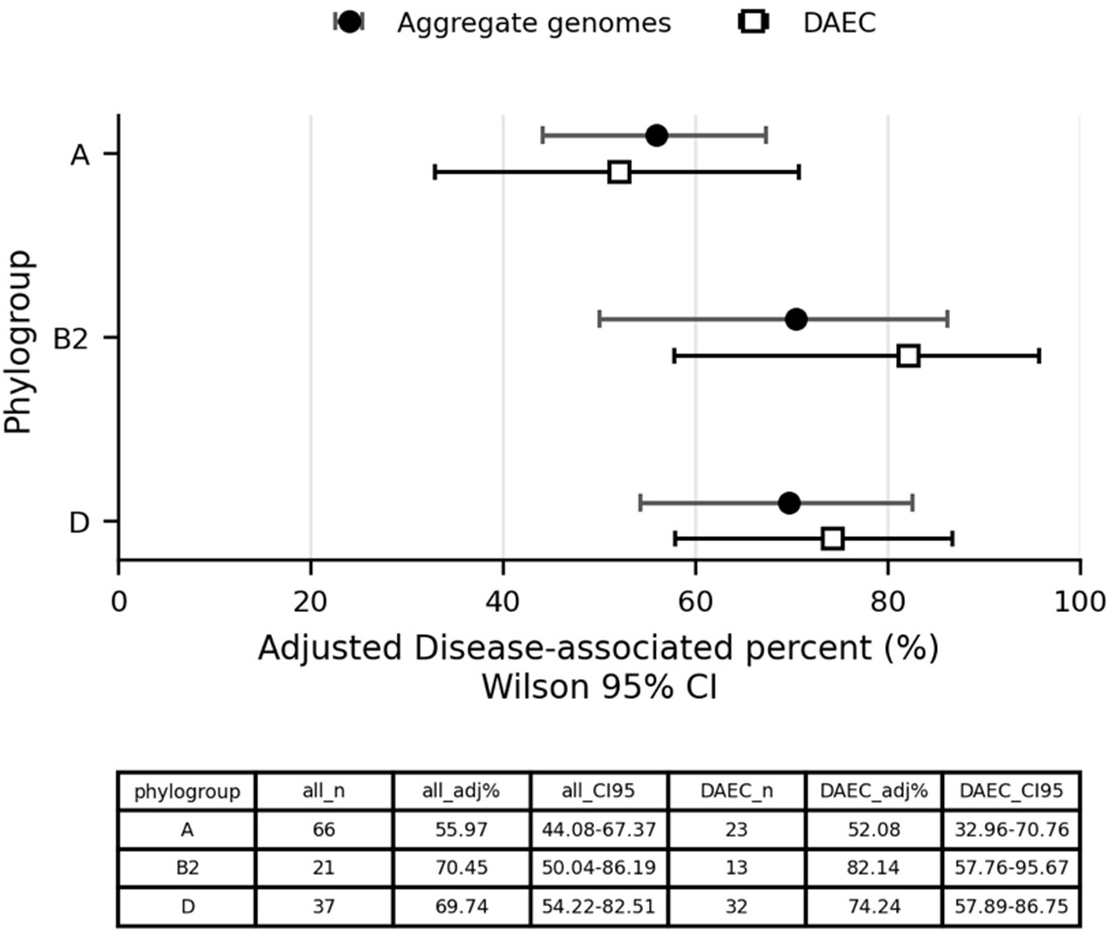
Disease association of *E. coli* phylogroups. Adjusted disease-association percentage by phylogroup. Black circles represent the aggregate trend across all genomes within each phylogroup, while open squares show the DAEC-specific trend. Error bars indicate Wilson 95% confidence intervals (CI). Disease association was defined as the proportion of diarrhea isolates relative to all retained genomes (diarrhea + asymptomatic) per phylogroup. Table shows the raw genome counts, adjusted disease-association percentages, and Wilson 95% CI for both the aggregate (all pathotypes) and DAEC-specific trends within each phylogroup.

**Supplementary Figure 7:**
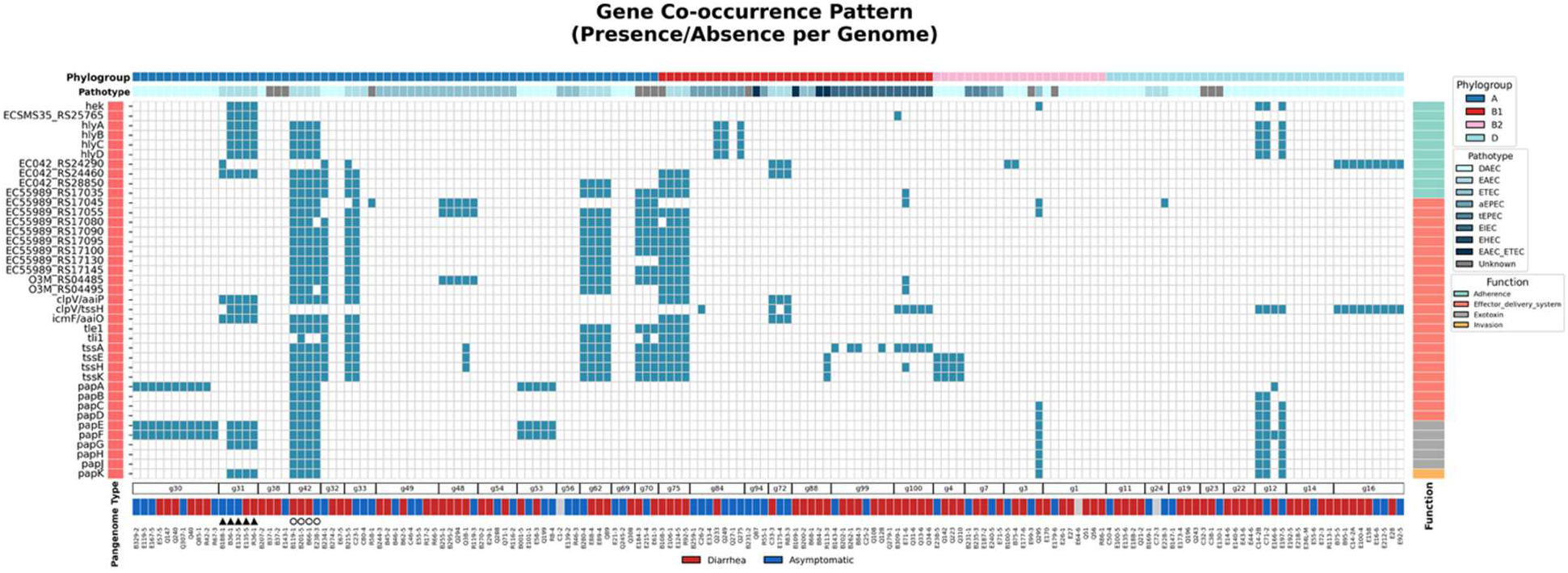
Distribution of rare virulence factors across phylogenetically ordered genomes reveals. gene clusters associated with diarrheal disease. The presence-absence heatmap displaying the union of rare virulence genes detected between genome members assigned to genomovars g31 (triangle) and g42 (circle) and compared across all genomes (x-axis) ordered according to their position in the core genome phylogeny (Figure 3). Genomes are annotated by phylogroup, pathotype, and clinical outcome status. The figure highlights contrasting rare gene profiles between the two closely related genomovars (with a genomovar mean genome-wide ANI = 99.23%, SD = 0.051), both belonging to the EAEC pathotype and assigned to ST-10 yet differing in disease prevalence. Cluster g42 predominantly carries a suite of putative effector delivery system components (genes with EC55989 prefix) and type VI secretion system genes (tssA, tssE, tssH, and tssK) that are absent in g31. The distribution of this gene union across additional genomovars reveals that g62, g70, and g75 (all assigned EAEC) also harbor these g42-associated genes and show elevated proportions of disease-associated isolates, while g31 and g24 lack these genes and exhibit lower disease association.

